# Weight initialization shapes task organization in recurrent neural networks

**DOI:** 10.64898/2026.08.08.743683

**Authors:** Renate Krause, Valerio Mante

## Abstract

Flexibly recombining computational modules is essential for biological and artificial neural networks to rapidly adapt to changing environments. This requires modules to be shared across tasks rather than rigidly segregated, yet what determines this organization remains unknown. Previous work suggests that weight initialization shapes whether networks learn task-specific or generic representations, but it is unclear whether this extends to recurrent networks and, more importantly, to network connectivity. Here, we systematically vary the initial weight variance of recurrent neural networks and study them using a framework that allows us to identify the functionally relevant connectivity subspaces for each computational module. We find that networks with low initial weight variance converge to solutions in which different subtasks rely on largely overlapping weight subspaces, whereas high-variance networks implement subtasks in higher-dimensional, more segregated weight subspaces. Our results also provide mechanistic insights with implications for interpreting biological neural circuits and for designing efficient recurrent architectures.

---

To rapidly adapt to diverse environmental conditions, biological and artificial neural networks must be able to flexibly reuse computational modules learned in previous contexts. However, such systems could implement any computational module either in a flexible, reusable fashion or as rigid, segregated components. Flexible and modular organization is initially more expensive to build but becomes more efficient over time, as learned modules can be reused across different settings. In contrast, rigid and segregated implementations are cheaper to build but can lead to inefficiency and redundancy when similar modules must be implemented multiple times. Hence, both types of organization have their own advantages, and understanding which factors determine where a network is placed on this spectrum remains an important open question.

Recent work has shown that biological and artificial neural networks can implement computational modules in a flexible and reusable manner. In the prefrontal cortex of rhesus macaques, Tafazoli et al. (*1*) showed that different neural subspaces are reused across similar tasks. Similarly, recurrent neural networks have been shown to flexibly reuse dynamical motifs and subspaces across tasks (*2, 3*), and comparable modular reuse has been observed in transformer-like architectures (*4*) or in feedforward networks using natural language instructions (*5*). However, these studies have focused on modularity at the level of network activity. It remains unclear how this organization manifests at the level of network connectivity and, importantly, what defines where on the spectrum between flexible, shared, and rigid, segregated task implementation a network lands.

We study this in recurrent neural networks (RNNs), as previous work has established direct links between different forms of recurrent connectivity and the resulting computational dynamics (hand-designed: (*6–8*); random networks: (*9–11*); constrained architectures: (*12–14*)). The concept of *low-rank RNNs* has proven especially valuable (*13*). Such networks can be explicitly constructed or trained to use a low-rank connectivity, enabling reverse engineering to provide insight into neural computation and population dynamics (*13, 15–17*). Even in networks trained without explicit low-rank constraints, connectivity can often be reduced to a low-rank approximation, as only a fraction of the weight subspaces is required to recover the original performance (*18*). This suggests that only a small subspace of the full connectivity is functionally relevant, motivating a framework to systematically identify such subspaces.

Previous work has shown that network solutions depend heavily on the initial conditions, specifically the initial variance of network weights, rather than on the learning algorithm itself (***?****, 19–22*). It has focused on feedforward architectures, describing learning as occurring in two distinct regimes: the *rich* (low initial weight variance) and the *lazy* (high initial weight variance) learning regimes. Rich learning typically shows slower learning, larger parameter updates, and more task-specific feature learning. In contrast, the lazy regime is associated with faster learning and less task-specific, higher-dimensional representations. Other structural factors (e.g., activation function (*2*), output weight initialization (*23*)) have also been shown to influence the solutions to which the networks converge. However, none of these factors has been directly linked to whether tasks are organized in a shared or segregated fashion at the level of recurrent connectivity, leaving open what controls this organization.

In this work, we study how functional modules are organized in the connectivity of RNNs and what controls this organization. We show that the initial variance of recurrent weights is a crucial factor shaping the dimensionality and organization of the functionally relevant connectivity. To this end, we apply the framework of operative dimensions (*18*) to identify task-relevant subspaces within the recurrent weight matrix and analyze how these operative subspaces differ across learning regimes. Operative dimensions reveal that networks trained in the rich regime develop lower-dimensional, more shared subtask representations, whereas lazy networks show higher-dimensional, more segregated ones. Our analyses also reveal a potential mechanism: these distinct organizations arise because networks in the rich regime tailor their weight updates more effectively, focusing adjustments primarily within the functionally relevant subspace of network connectivity rather than uniformly across the entire weight matrix. Together, these results show that weight initialization is an important factor in determining how computational modules are organized within recurrent connectivity, with implications for both interpreting biological neural circuits and designing efficient recurrent architectures.

## Network setup

We perform our analyses on vanilla RNNs using the following standard RNN equation:

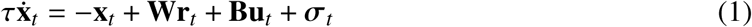

The *network inputs* **u***_t_* ∈ R*^U^* are task-dependent and time-varying and sent to the *N* = 100 hidden units with input weights **B** ∈ R*^N^*^×*U*^. The *recurrent weight matrix* of the hidden units is given as **W** ∈ R*^N^*^×*N*^. **x***_t_* ∈ R*^N^* are the linear activities of the hidden units at time *t* with **r***_t_* = *tan*ℎ(**x***_t_*) (tanh) or **r***_t_* = *max*(0, **x***_t_*) (ReLU). The hidden units are noisy, where each element of ***σ****_t_* is drawn from a Normal distribution N ( *μ* = 0, *σ* = 0.1). *τ* ∈ R is the time constant (*τ* = 10 *ms*, *dt* = 1*ms*).

The *network output* **z***_t_* ∈ R*^Z^* is defined as:

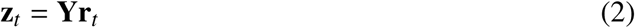

with output readout weights **Y** ∈ R*^Z^*^×^*^N^*

For any given condition, the cost is defined as:

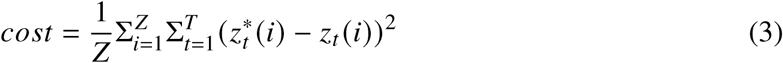

where 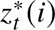 is the desired output. All network weights are trained to minimize the summed costs across all input conditions. Only the *network inputs* **u***_t_* and the randomly drawn ***σ****_t_* vary across input conditions. All network weights (**B**, **W**, **Y**) are trained either with backpropagation through time (BPTT) (*24*) with the Adam optimizer (*25*), (*lr* = 1*e* − 4) or Hessian-free optimization (HF) (*26*) to minimize the cost (Eq. 3). All networks are trained without any regularization terms.

## Tasks

The RNNs are trained separately on two different tasks. The first task is context-dependent integration (CTXT, Fig. 1a,b) (*27*). The RNN receives two noisy, sensory inputs and two static, context inputs (*U* = 4). The network is trained to select one of the two sensory inputs (depending on the currently active context input; i.e., select input sensory_i_ in context_i_) and integrate it over time (Fig. 1a,b).

**Figure 1:**
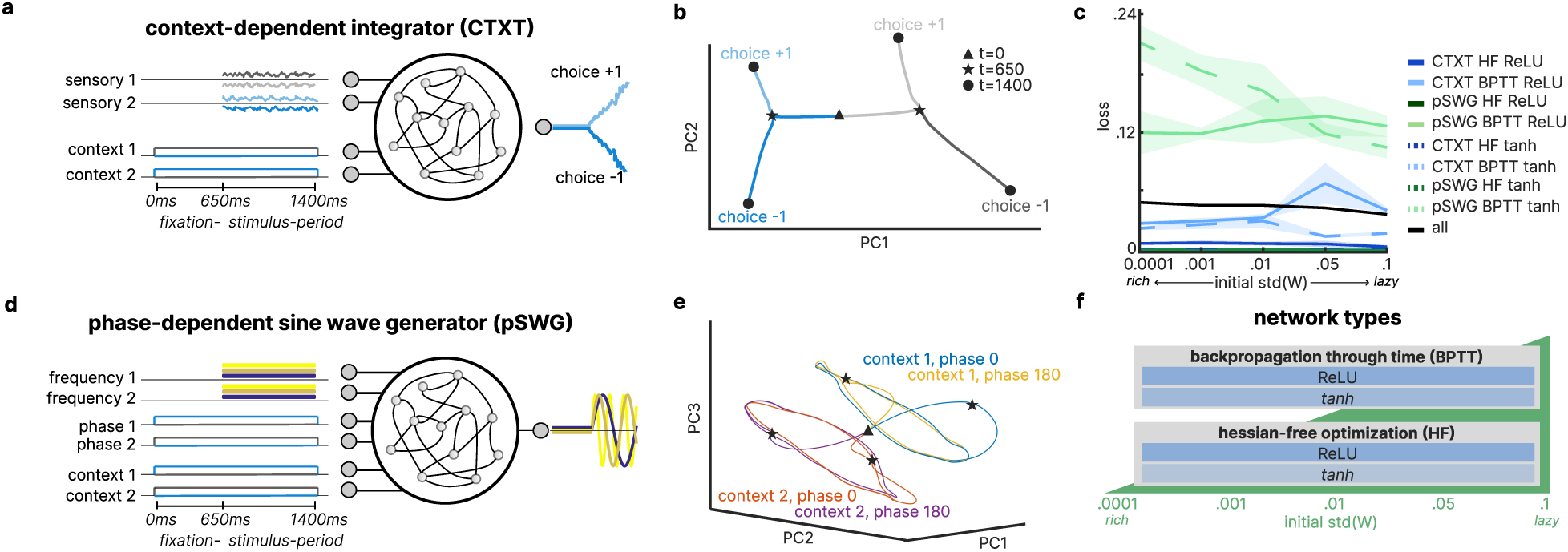
Network setups. (**a**) Network architecture for context-dependent integration (CTXT). (**b**) Low-dimensional projection of condition average trajectories in CTXT (1 representative example network, *N* (0, [1*e*−4])). (**c**) Loss on test set at the end of training for eight different network families over varying weight initializations (shaded area: median absolute deviation (*mad*) over all networks per network setting). (**d**) Network architecture for phase-dependent sine wave generation (pSWG). (**e**) Low-dimensional projection of condition average trajectories in pSWG (1 representative example network, *N* (0, [1*e* − 4])). (**f**) Illustration of studied network settings.

The second task of phase-dependent sine wave generation (pSWG, Fig. 1d,e) is an extension of the traditional sine wave generation task (*28*). The RNN is trained to output a sine wave at a specified frequency and phase (0 or 180 degrees). It receives two static inputs that determine the frequency of the generated sine wave, two static inputs that determine the phase of the generated sine wave (0 or 180 degrees), and two static inputs that determine which of the two frequencies or phases the network should follow (*U* = 6) (Fig. 1d,e). The pSWG task was designed to complement CTXT by engaging a qualitatively different dynamical motif: whereas CTXT typically yields a static attractor (line attractor, (*27, 29*)), sine wave generation relies on dynamic attractors (*28*). Both tasks share the overall structure of modular neuroscience paradigms, with distinct input conditions defining subtasks.

Both tasks consist of two *task periods*: *fixation period* (fix) and *stimulus period* (stim). In the first period, only the context inputs are activated, and the network is instructed to keep **z***_t_* = 0 (*fixation period*: = 650 *ms*). In the second period, the stimulus inputs are switched on, and the network is asked to generate the desired non-zero outputs as described above (*stimulus period*: = 750 *ms*).

## Weight initializations

To study the *rich* and *lazy* learning regimes, the recurrent weights **W** are initialized with five different variances: *N* (0, [1*e* − 4, 1*e* − 3, 0.01, 0.05, 0.1]). Training failed to converge for the majority of networks initialized with variances larger than 0.1. Consistent with previous literature (*20–22, 30*), we refer to networks with small initial variance in **W** as being trained in the *rich* regime and to networks with large initial variance in **W** as being trained in the *lazy* regime. The input and output weights (**B**, **Y**) are randomly initialized with *N* (0, 0.1).

In summary, we trained RNNs on two tasks (CTXT, pSWG), using two activation functions (tanh, ReLU), two gradient-based optimization methods (BPTT, HF), and five different initial variances for the recurrent weights (Fig. 1f). Although the different network families reach slightly different final performance levels, all networks show clear convergence of both training and test loss during training (Fig. 1c). To ensure comparability across families, all networks were trained with the same number of training samples and training epochs (120 epochs; see Supplementary Materials for details).

## Linking structure to function with operative dimensions

Recurrent connectivity matrices (**W**) are typically high-dimensional (*18, 30*) and Fig. AS1, unless explicitly constrained by regularization, making them difficult to interpret. Nevertheless, analyzing the network connectivity is essential: any changes in network performance are ultimately implemented through updates to the connectivity.

To systematically link network function to network connectivity in recurrent neural networks, we use the framework of operative dimensions (*18*). Operative dimensions identify the functionally relevant subspaces within the recurrent weight matrix **W** (Fig. 2a), thereby providing a direct link between structure and function. Intuitively, operative dimensions are defined through an iterative optimization procedure that identifies directions in the recurrent connectivity with maximal impact on the network function. Here, impact refers to changes to the local dynamics of the network at specific states **x***_t_*. In other words, operative dimensions correspond to perturbations of the recurrent weights that maximally disrupt the local flow field (see formal definition in Supplementary Methods). By successively identifying such directions in the weight matrix, one obtains a set of dimensions (in weight space) ranked by their functional relevance for the network function.

**Figure 2:**
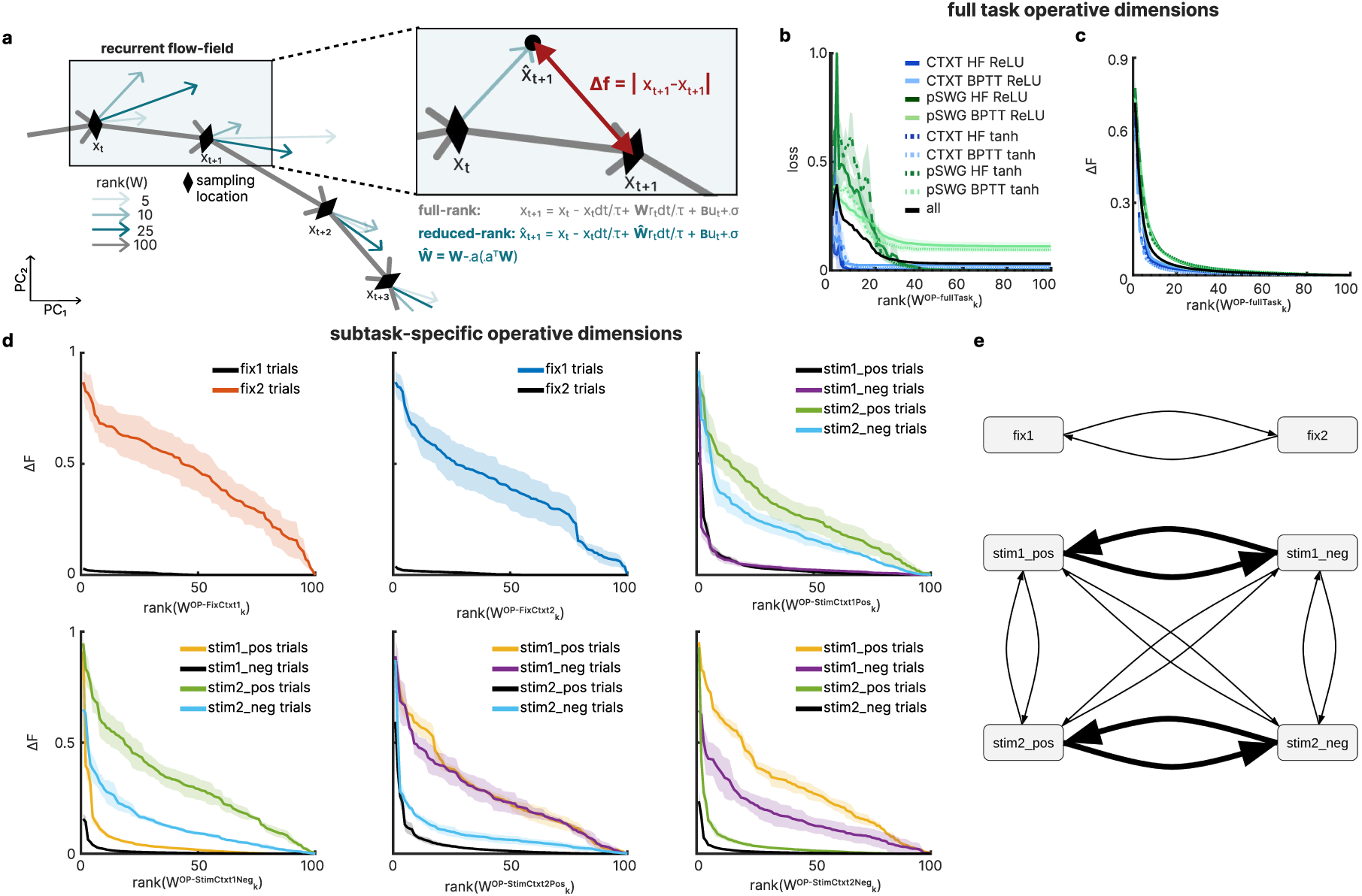
Operative dimensions. (**a**) Illustration of operative dimensions showing the recurrent flow field along the network trajectory. Colored arrows show dynamics generated by the full-rank recurrent weight matrix (**W**) and several reduced-rank approximations (**Ŵ**, ***σ****_t_* = 0, colors per legend). Local operative dimensions maximize Δ *f*. (**b**) Test loss / (**c**) Δ*F* of networks with reduced-rank weight matrix 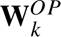 for rank *k* = 1: *N* (Eq. 4) of eight different network types, as illustrative example here for *σW*_0_=0.1. (**d**) Δ*F* per subtask for networks with a reduced-rank weight matrix consisting of the first *k* subtask-specific operative dimensions. Here is an illustrative example of a network trained on context-dependent integration (CTXT, HF, tanh) showing that the subtask-specific operative dimensions are only partially shared even between closely related subtasks (shaded area is *mad* over 20 noisy trials per subtask type). (**e**) Graphical illustration of the relationship between the different subtasks, approximated based on similarity in (**d**). Line thickness is proportional to the similarity of the operative dimension subspace.

## Low-dimensional weight subspace is sufficient for original task performance

Crucially, reconstructing the weight matrix using only a small number of operative dimensions preserves the network’s original performance (Fig. 2b,c). Hence, most of the seemingly high-dimensional network connectivity (Fig. AS1) is not functionally relevant to the task nor directly involved in generating the trained network function (Fig. 2b,c, see also previous results for more complex task (sequential MNIST) (*18*)).

To reveal that the recurrent weight matrix is functionally low-rank, we iteratively removed operative dimensions from **W** to construct reduced-rank approximations of **W**, and tested the network’s performance. For column dimensions, the reduced-rank approximations 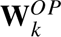 are:

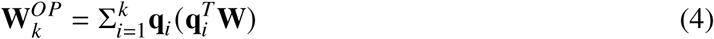

First, we compare the network’s performance, as defined in Eq. 3 (Fig. 2b) over 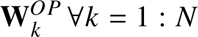. Additionally, we compare the network trajectories generated using 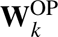 with those of the original, full-rank network **W** (Fig. 2c)). The difference in network trajectories is referred to as Δ*F*. It is quantified as the average change in its local dynamics (Δ*f*) at network states of the original, full-rank network, normalized by the average step size of the network activity (|**x***_t_*_+1_ − **x***_t_* |∀*t* = 1: *T*). This normalization accounts for differences in activity magnitude across hyperparameter settings.

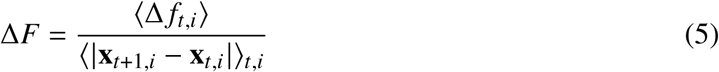

In this work, we averaged Δ *f* at time steps *t* (*t* = 1: *T*; *T* = 1400) in random trials *i* (trials equally distributed across input condition; *i* = 1: 40). A low-dimensional subspace of the originally high-dimensional recurrent weight matrix is sufficient to achieve both the network’s original performance and dynamics (Fig. AS1, Fig. 2b,c). Hence, task-relevant computations are limited to a low-dimensional subspace of the overall weight space. The exact dimensionality of the subspace spanned by the operative dimensions is highly dependent on the specific criteria used to set the threshold. For CTXT < 20/100 dimensions are required to achieve approximately the original cost, for pSWG < 40/100 dimensions.

Consequently, we can restrict our analysis to this low-dimensional operative subspace of the original high-dimensional weight matrix to study the network’s functional organization. The remaining dimensions contribute little or nothing to the network dynamics. By focusing on these operative subspaces, we reduce connectivity complexity and effectively filter out functionally irrelevant components, enabling a more targeted analysis of how learning regimes shape recurrent structure. As we show below, restricting the connectivity analyses to the operative subspace is critical to identify some of the key differences between the rich and lazy regimes.

## Subtask-specific operative dimensions are distinct yet coherently structured

The framework of operative dimensions allows not only the extraction of the functionally relevant subspace for the entire task (see Supplementary Methods; Fig. 2b,c), but also for arbitrarily defined *subtasks* or even single time points. In this work, we focus on operative dimensions defined for individual subtasks, specified by their task periods (fixation period, stimulus period) and specific input conditions (*2, 31*). In total, we defined six subtasks for the context-dependent integration task and eight subtasks for the phase-dependent sine-wave generator task (Table 1). For each of them, we can then define a specific set of operative dimensions.

**Table 1:** Overview of subtasks for each task. The table lists the abbreviations used for each subtask together with a detailed description of the corresponding task, task period, and specific input conditions defining each subtask.

| subtask name | task | task period | input condition |
| --- | --- | --- | --- |
| fix1 | CTXT | fix | $\mathbf{u}_t = [0, 0, 1, 0]$ |
| fix2 | CTXT | fix | $\mathbf{u}_t = [0, 0, 0, 1]$ |
| stim1_pos | CTXT | stim | $\mathbf{u}_t = [i, j, 1, 0] \forall i > 0, j \in \mathbb{R}$ |
| stim1_neg | CTXT | stim | $\mathbf{u}_t = [i, j, 1, 0] \forall i < 0, j \in \mathbb{R}$ |
| stim2_pos | CTXT | stim | $\mathbf{u}_t = [i, j, 0, 1] \forall j > 0, i \in \mathbb{R}$ |
| stim2_neg | CTXT | stim | $\mathbf{u}_t = [i, j, 0, 1] \forall j < 0, i \in \mathbb{R}$ |
| fix1_ph0 | pSWG | fix | $\mathbf{u}_t = [0, 0, 1, 0, 1, 0]$ |
| fix1_ph180 | pSWG | fix | $\mathbf{u}_t = [0, 0, 0, 1, 1, 0]$ |
| fix2_ph0 | pSWG | fix | $\mathbf{u}_t = [0, 0, 1, 0, 0, 1]$ |
| fix2_ph180 | pSWG | fix | $\mathbf{u}_t = [0, 0, 0, 1, 0, 1]$ |
| stim1_ph0 | pSWG | stim | $\mathbf{u}_t = [i, j, 1, 0, 1, 0] \forall i, j > 0$ |
| stim1_ph180 | pSWG | stim | $\mathbf{u}_t = [i, j, 0, 1, 1, 0] \forall i, j > 0$ |
| stim2_ph0 | pSWG | stim | $\mathbf{u}_t = [i, j, 1, 0, 0, 1] \forall i, j > 0$ |
| stim2_ph180 | pSWG | stim | $\mathbf{u}_t = [i, j, 0, 1, 0, 1] \forall i, j > 0$ |

Comparing these subtask-specific operative dimensions provides insight into how different subtasks are organized within the network connectivity. To illustrate their relationships, we evaluated how well operative dimensions derived from one subtask generalize to others (assessed using Δ*F*, Eq. 5), shown here for one representative network trained on CTXT in the lazy regime (*σW*_0_=0.1) (Fig. 2d). When the network connectivity is constrained to the operative dimensions derived for its own fixation period in a given context (e.g., *OP* for fix1 to run trials of fix1), fewer than 20 dimensions are sufficient to recover approximately the original network trajectories (Δ*F* ≈ 0). In contrast, when operative dimensions from the fixation period of the other context are used (e.g., *OP* for fix2 to run trials of fix1), nearly the full set of 100 dimensions is required to achieve a comparable Δ*F* (Fig. 2d), indicating that these subspaces differ substantially.

A similar pattern emerges in the stimulus period: weight subspaces are more similar between subtasks of the same context (e.g., stim1 pos vs stim1 neg) than between subtasks sharing the same choice across contexts (e.g., stim1 pos vs stim2 pos), as reflected in lower or higher Δ*F* for the same *rank* (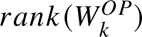) (Fig. 2d). Together, these results demonstrate that each subtask relies on a low-dimensional subspace of the weight matrix. These subspaces are only partially shared and not interchangeable, yet they exhibit a structured overlap that reflects the underlying task relationships (Fig. 2e).

## Task organization across learning regimes

In the following, we turn to the question of how computational modules are organized and represented in the RNNs across different learning regimes.

We first checked whether our network setup reproduces key qualitative differences between rich and lazy learning regimes reported in previous studies, which were mainly conducted on feedforward networks (see (*21*) for an informative review). As in feedforward architectures, our lazy networks learn faster, with the loss decaying and converging after fewer training epochs. The loss in lazy networks also decreases more smoothly, without intermediate plateaus (Fig. 3a, e.g., (*20, 30, 32*)). In addition, networks initialized with higher variance in **W** tend to maintain this elevated variance throughout training (Fig.A S8).

**Figure 3:**
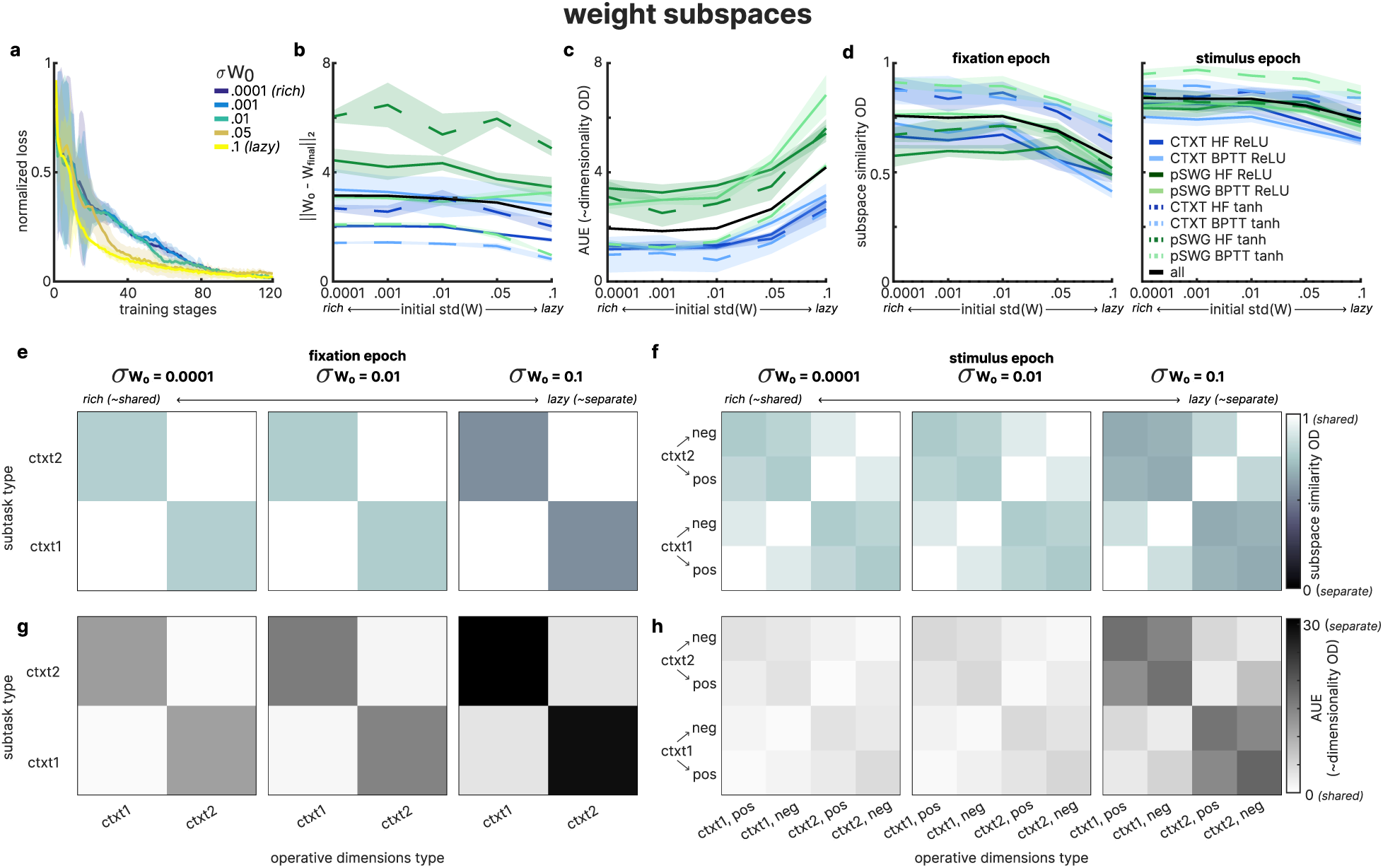
Task representations in network connectivities. (**a**) Normalized test loss for example network type (CTXT, HF, tanh) over training epochs for varying weight initializations *σW*_0_. (**b**) Absolute difference between **W** before and after training (L2-norm) in eight different network types over varying weight initializations *σW*_0_. (**c**) Area Under Error (*AUE*) for eight different network types over varying weight initializations *σW*_0_. The *AUE* increases with larger initial weight variance, indicating an increase in the dimensionality of the functionally relevant weight subspace. (**d**) Average, pairwise similarity of all subtask-specific operative dimensions per network in eight different network types over varying weight initializations *σW*_0_. (**e**) Pairwise similarity of subtask-specific operative dimensions of fixation period in CTXT (averaged over networks) over varying weight initializations *σW*_0_. (**f**) same as (**e**) for stimulus period. (**g**) *AUE* of subtask-specific operative dimensions of fixation period in CTXT (averaged over networks) over varying weight initializations *σW*_0_. *AUE* is calculated for each subtask type (y-axis) using all subtask-specific operative dimension types (x-axis). (**f**) same as (**g**) for stimulus period. (**a**)-(**d**) shaded area: median absolute deviation (*mad*) over all networks per *σW*_0_.

However, in contrast to previous findings in feedforward ANNs (*20, 32*), lazy networks here do not exhibit a consistently smaller overall change in the recurrent weight matrix **W** over the course of training. Only a slight trend can be observed in that direction (Fig. 3b). Since previous studies have shown that only a fraction of the recurrent weight matrix **W** contributes meaningfully to network function (*18*), we next analyze the network connectivity using the framework of operative dimensions to isolate the functionally relevant subspaces for further analysis.

## Lazy learning regime yields higher-dimensional task implementations

In a first step, we assessed the dimensionality of these functionally relevant weight subspaces for the entire task. To this end, we obtained operative dimensions for all network types and assessed how many of these dimensions in the weight matrix were required for the network to perform the task. Specifically, we sequentially removed operative dimensions from the weight matrix (as explained above, see (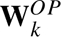 in Eq. 4) and measured the resulting performance (Δ*F*, Eq. 5) of the reduced-rank networks. To summarize the resulting error curves for Δ*F*, we compute the *area under the error curve* (*AUE*, Fig. 3c) for each network. A larger *AUE* indicates a higher-dimensional functionally relevant subspace, and a smaller *AUE* indicates lower dimensionality.

We find that networks with smaller initial weight variance (rich regime) consistently show lower *AUE*, indicating that they rely on lower-dimensional functional subspaces to solve the same task. By contrast, networks with larger initial weight variance (lazy regime) exhibit systematically higher *AUE* values, with the dimensionality of their functional subspaces gradually increasing as the initialization variance increases. This observation holds across all tasks, training methods, and activation functions, averaged over networks (Fig. 3c).

## Rich learning regime yields more shared task implementations

At a finer level, we can study the organization of subtask-specific weight subspaces (see Table 1). To this end, we measure the similarity of the subspaces spanned by the subtask-specific operative dimensions using the concept of principal vectors (see Materials and Methods), which intuitively reflects the average overlap between two subspaces.

Applying this similarity measure to subtask-specific operative dimensions, we observe that subtasks of rich networks are more similar to one another than those of lazy networks. As illustrated in Fig. 3d-f, networks trained in rich and lazy regimes exhibit a qualitatively similar organization of subtasks (see Fig. AS3 for pSWG). However, as we move from the rich to the lazy regime, the subspaces become progressively less similar, showing greater separation.

Additionally, we can assess the similarity of subtask-specific operative dimensions at the level of network function (see comparison of measures in Fig. AS6). To this end, we take the subtask-specific operative dimensions identified for one subtask and test the network performance when applying them in a different subtask. We again quantify performance using the area under the error curve (*AUE*). This provides a measure proportional to the dimensionality of the operative subspace that the network requires to solve the task. Specifically, a small *AUE* indicates that the weight subspaces recruited in one subtask are mostly shared with those of the other subtask. A large *AUE* indicates that the recruited subspaces are clearly distinct.

Using this similarity measure, we observed the same trend as in the subspace similarity analysis: rich networks exhibit substantial sharing of operative subspaces across subtasks, whereas in lazy networks, the degree of sharing decreases progressively with the variance of the initial weights (Fig. 3g,h). Interestingly, weight subspaces are more shared across subtasks of the same context (e.g., *ctxt*1 *pos*, *ctxt*1 *neg*) than across subtasks sharing the same choice (e.g., *ctxt*1 *pos*, *ctxt*2 *pos*). This qualitative organization is consistent across both rich and lazy learning regimes (Fig. 3g,h).

Notably, this organization is not simply inherited from initialization, as subtask subspaces are organized very differently before training than after. At initialization, rich networks show less shared operative subspaces than lazy networks (Fig. AS4a). In rich networks, early training quickly makes these subspaces more similar to each other (Fig. AS4b). In contrast, lazy networks show very little change in subspace similarity over early training (Fig. AS4b). Across learning, subtasks within the same context remain more similar to one another in both regimes. Overall, this suggests that the final subtask organization is mainly shaped by the learning dynamics induced by the initialization, rather than simply reflecting the initial organizational structure.

## Differences in task implementations are reflected in network activities

We observe similar patterns when analyzing the network activity itself. First, we find that lazy networks generally exhibit higher-dimensional dynamics after training than rich networks (Fig. 4a-b). In addition, the activity subspaces associated with different subtasks become progressively less shared as networks move further into the lazy regime. To quantify these observations, we compared the similarity of the subspace spanned by the first *k* principal components (PCs) of the network activity across subtasks *S_PCA_*_,*PC*_*_B_*. In rich networks, the subtask-specific activity subspaces are relatively well aligned within each network. However, when averaging across all network types, the same trend emerges as for the weight-subspace analysis: subtask-specific subspaces are more strongly shared in rich networks and become increasingly distinct in the lazy regime (Fig. 4c-e, see Fig. AS5 for pSWG). The similarity between operative dimensions and the PCs of network activity observed here is consistent with previous findings (*18*), but it is not guaranteed in general (see Discussion).

**Figure 4:**
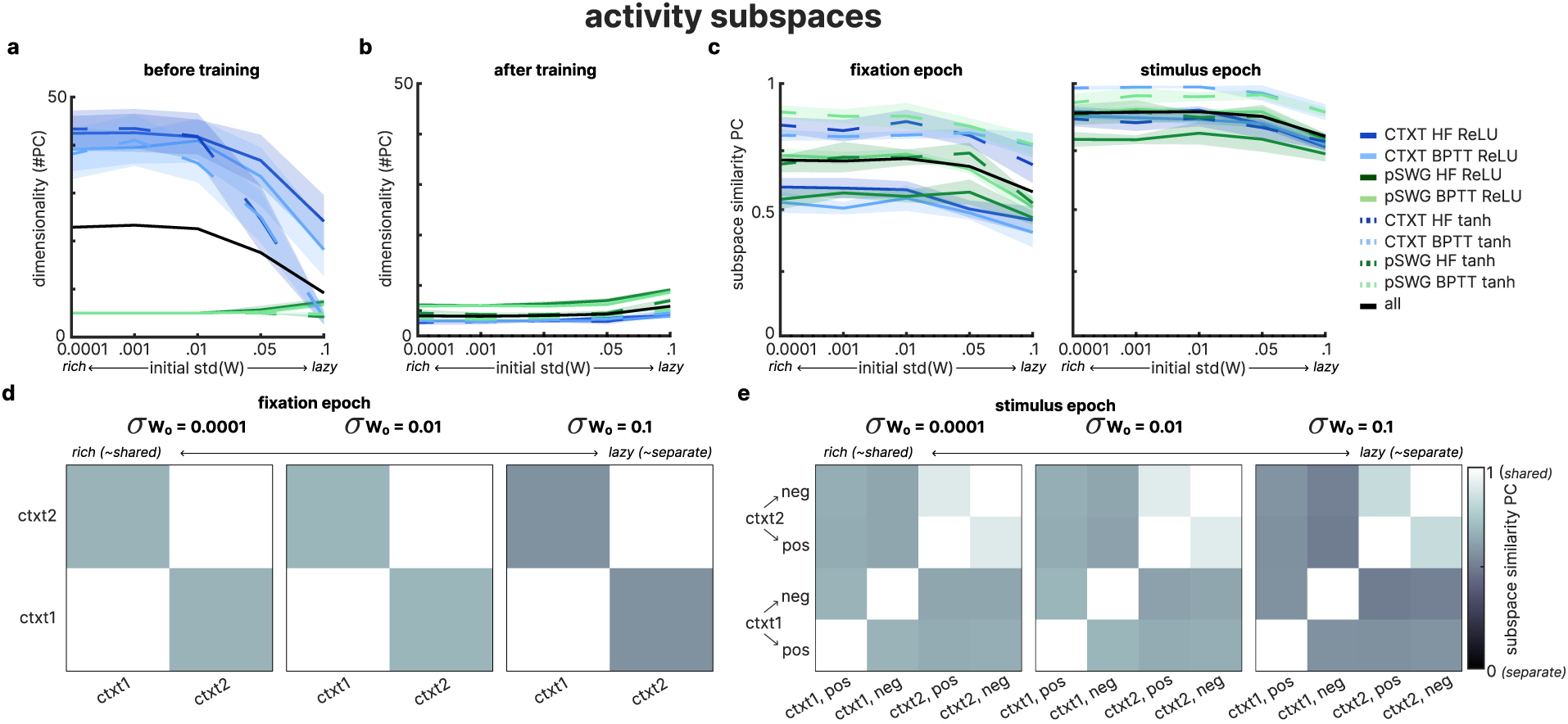
Task representations in network activities. (**a**)/(**b**) Dimensionality of network activities before (**a**) and after (**b**) training, measured as the number of PCs required to explain 95% of the variance in network activities (shown for all networks per network type). (**c**) Average pairwise similarity of subtask-specific network activity subspaces (first *k* PCs) per network in eight different network types over varying weight initializations *σW*_0_ during fixation and stimulus period. (**d**) Pairwise similarity of subtask-specific PCs of fixation period in CTXT (averaged over all networks) over varying weight initializations *σW*_0_. (**e**) same as (**d**) for stimulus period. (**a**)-(**c**) *mad* over all networks per *σW*_0_.

Overall, our analyses reveal consistent patterns across subtasks when comparing weight-based subspaces (operative dimensions) and activity-based subspaces (principal components). In both cases, networks trained in the rich regime rely on lower-dimensional subspaces that are strongly shared across subtasks. In contrast, networks in the lazy regime exhibit higher-dimensional solutions with subspaces that are progressively less overlapping across subtasks. Thus, both the connectivity-level analysis using operative dimensions and the activity-level analysis using PCA converge on the same conclusion: rich networks implement more compact, shared functional subspaces, while lazy networks rely on larger, more segregated subspaces.

## Learning trajectories across task organization

Given that the networks converge to qualitatively different solutions after training, we next examine the learning dynamics to understand how these distinct solutions emerge across the two learning regimes. The difference in weight variance between the two regimes is most pronounced early in training. Rich networks gradually increase the variance of the recurrent weights over training, whereas lazy networks show only a slight increase or remain relatively stable (Fig. AS8). Therefore, in the following analyses, we focus primarily on the early training phase (epochs 1–20), during which the distinction between the two learning regimes is strongest. Notably, we find that the two regimes differ both in where weight updates are applied and in how these updates reshape the operative subspaces.

## Rich networks perform more targeted recurrent weight updates

Most prominently, rich networks exhibit more *targeted* weight updates (Fig. 5a), meaning a larger fraction of the recurrent weight changes occurs within the functionally relevant subspace of **W**. We measured the proportion of total recurrent weight update that lies within the operative subspace of 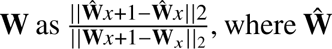 is reconstructed using exclusively the first five operative dimensions of iteration *x* (see Eq. 4, *k* = 5; consistent across subspace dimensionality Fig. AS9a). This measure yields values in the range [0, 1], where 0 indicates that all weight updates were performed along dimensions outside the functionally relevant weight subspace, and 1 indicates that all weight updates were performed within it. Across all architectures, rich networks show more specific learning: a larger proportion of their recurrent weight updates lies within the functionally relevant subspace of **W**, whereas lazy networks distribute weight updates more broadly. Interestingly, lazy networks still exhibit faster reductions in loss despite their seemingly less specific weight updates.

**Figure 5:**
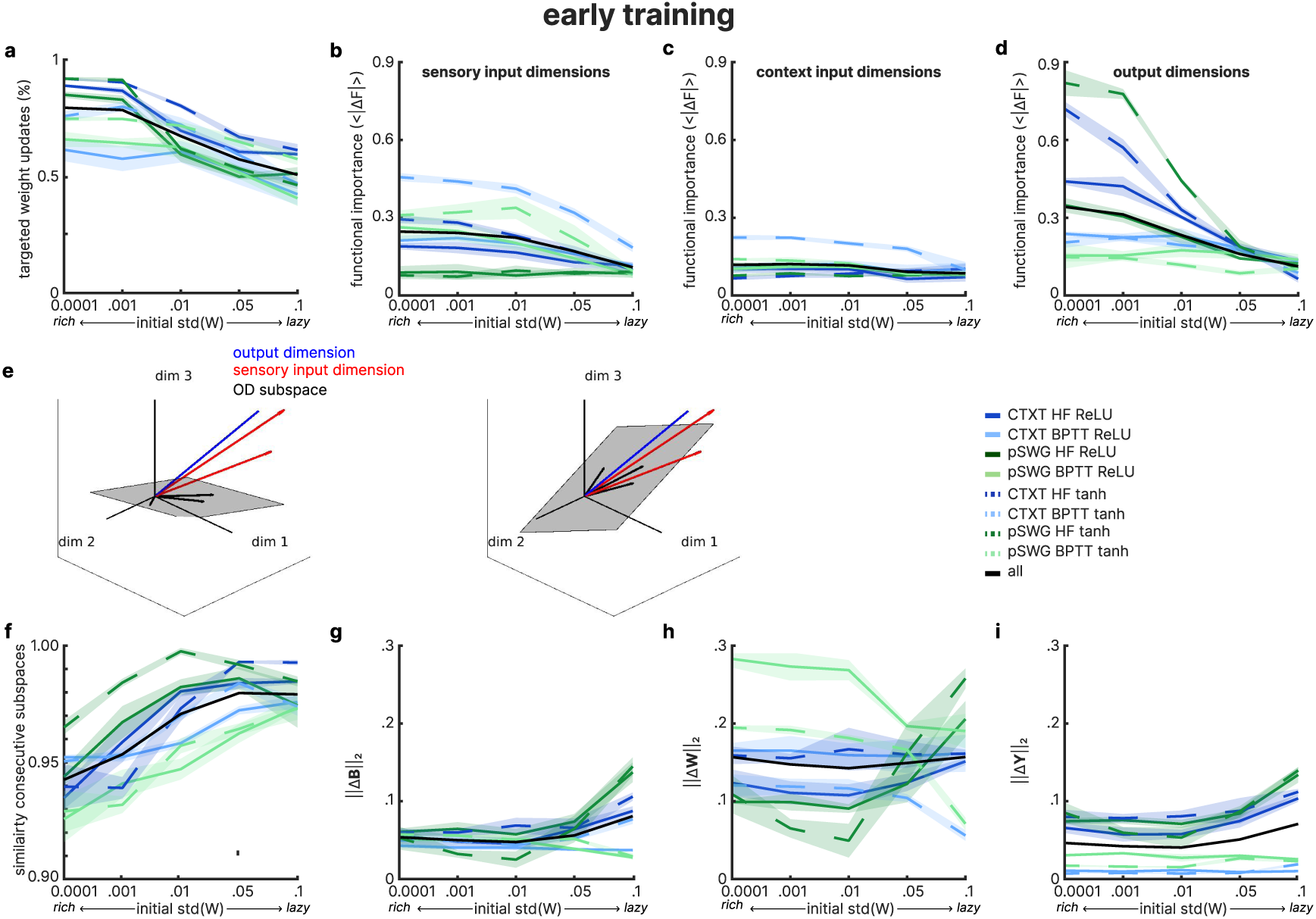
Differences across learning regimes early in training. (**a**) Average fraction of weight updates performed in the functionally relevant subspace of **W** (averaged over all weight updates of training iterations 1-20). (**b**) Δ*F* of removing the sensory input dimensions (**B**_1,2_) (averaged over training iterations 1-20). (**c**) Δ*F* of removing the context input dimensions (**B**_3−6_) (averaged over training iterations 1-20). (**d**) Δ*F* of removing the output dimension (**Y**) (averaged over training iterations 1-20). (**e**) Schematic illustration of possible alignments between input-dimensions (red), output-dimensions (blue), and operative dimension subspace (black). (**f**) Similarity (*S*) of the functionally relevant subspace of **W** between adjacent training epochs (averaged over training iterations 1-20). (**g**) Total difference in input weights **B** over training iteration 1-20. (**h**) Total difference in recurrent weights **W** over training iteration 1-20. (**i**) Total difference in output weights **Y** over training iteration 1-20. (**a**)-(**h**) shaded area: median absolute deviation (*mad*) over all networks per *σW*_0_.

A complementary analysis provides insight into why rich networks perform more targeted updates. Specifically, we assessed the functional importance of the network’s input and output dimensions (given by **B** and **Y**) by removing these dimensions from **W** and quantifying their impact on the network dynamics via Δ*F* (Eq. 5). Intuitively, this measures how strongly input and output dimensions contribute to the functionally relevant subspace (Fig. 5e). Our results indicate that dimensions aligned with the *sensory* inputs (**B**_1−2_) and the output weights **Y** have a stronger impact on network dynamics in rich than in lazy networks (epoch 1-20, Fig. 5b,d). A weaker but consistent pattern is observed for the *context* input dimensions (**B**_3−6_) (Fig. 5c). Thus, in the rich learning regime, input and output dimensions are more strongly integrated into the functionally relevant recurrent subspace. Crucially, this provides a potential mechanism for the more targeted gradient-based weight updates in rich networks. This interpretation is supported by theoretical analyses of how gradient-based learning operates (*30, 33, 34*) (see discussion). However, if alignment with the operative subspace plays such a central role, how does this subspace itself evolve over training?

## Rich networks rearrange their functionally relevant subspaces more than lazy

To address the evolution of the operative subspace during early training in greater detail, we quantified how its location changes during weight updates. We computed the average subspace similarity *S* (see Eq. S8) between subspaces of consecutive training epochs (epoch 1–20). We find that the operative subspace in rich networks shifts substantially during early learning, whereas in lazy networks it remains much closer to the subspace defined at initialization (Fig. 5f; consistent across subspace dimensionality Fig. AS9b). Thus, rich networks not only concentrate their weight updates within the operative subspace but also actively reshape it as learning unfolds. For a detailed discussion on the relation between targeted weight updates and their impact on the operative subspace (see Materials and Methods).

A similar pattern emerges when examining the overall change in the functional weight subspace from before to after training (epochs 1–120). Learning alters the utilized weight subspace more strongly in rich than in lazy networks (see Fig. AS2 for development of *AUE* across training stages). Specifically, we compute the *AUE* for the initial network using the operative dimensions derived from the final network, and vice versa (Fig. AS7d–e). As shown in Fig. S5d-e, *AUE* values are larger for rich networks, indicating that operative dimensions of the initial network provide a poorer fit to the final network in the rich regime than in the lazy regime. In other words, the functionally relevant connectivity subspace undergoes more substantial reorganization in rich than in lazy networks over training.

Notably, lazy networks show only minor shifts in both the location and the structure of their operative subspace (Fig. 5a, f). Also, changes in the recurrent weights are comparable across regimes when assessed at the level of the full weight matrices (epochs 1–20; Fig. 5h; see Fig. AS7a-c for changes across the entire training). Nevertheless, lazy networks exhibit a faster and larger decrease in the loss (Eq. 3) during the early training epochs (Fig. 3a). Hence, they achieve rapid performance improvements with relatively stable recurrent connectivity, but stronger updates to the input and output weights than rich networks (Fig. 5g,i).

## Rich networks transiently reorganize subtasks during learning

To better understand how the functional organization of the subtasks emerges in detail, we examined how the relationships between subtask-specific operative subspaces change over training. For this analysis, we focused on one network trained on the context-dependent integration task (two representative networks trained on CTXT with HF; one ReLU, one tanh; Fig. 6a-b and c-d, respectively) and computed pairwise similarities among the operative dimensions of all subtasks at each training iteration (epochs 1-20). As a similarity measure, we used the subspace similarity (Eq. S8) of the subspace spanned by the first 5 operative dimensions. These similarities were then visualized both as a similarity matrix (Fig. 6a,c) and using multidimensional scaling (non-metric MDS) with 1 − *S* (Eq. S8) as the distance metric (Fig. 6b,d).

**Figure 6:**
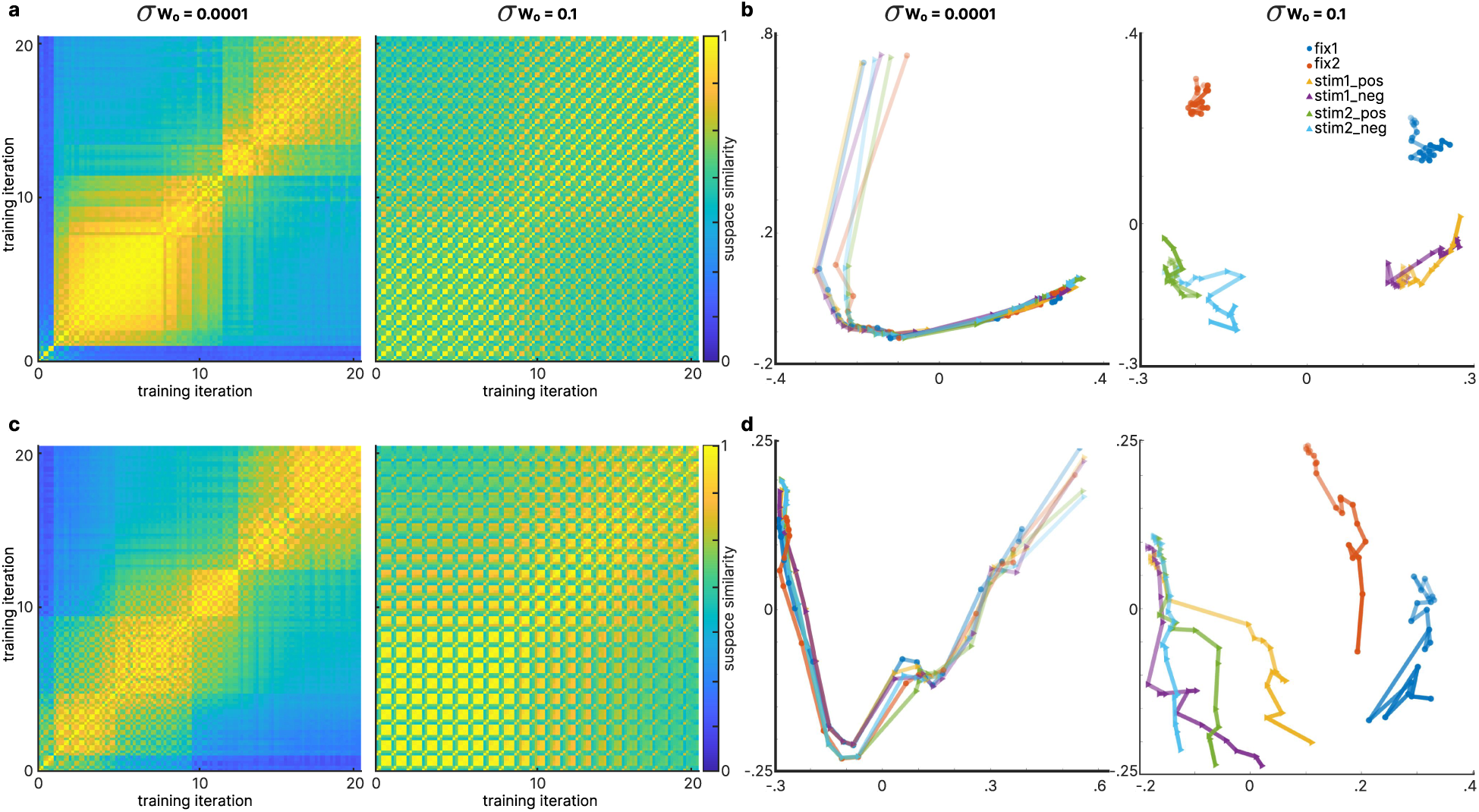
Subtask-specific operative dimensions over training. (**a**) Symmetric distance matrix (pairwise subspace similarity) over subtask-specific operative dimensions over training epoch 1-20 for networks with *σW*_0_=0.0001/0.1. The rows/columns are sorted as defined in the legend in (b) and repeated for all shown (20) training epochs. (**b**) MDS on pairwise subspace similarity between subtask-specific operative dimensions for networks with *σW*_0_=0.0001/0.1 (MDS stress=0.103/0.165). Dot color indicates subtask type. Color saturation indicates training epoch (light=epoch 0, dark=epoch 20). Both plots show data from one example network trained on CTXT, using HF with a ReLU activation function. (**c**)-(**d**) Same as (**a**)-(**b**) but for example network trained on CTXT, using HF with tanh-activation function (MDS stress=0.089/0.214).

The evolution of the subtask organization differs markedly between the rich and lazy learning regimes. In the rich regime, the relationships between subtasks change substantially during early training. Initially, the subtasks get grouped by context, then this grouping dissolves, and the operative subspaces of all subtasks temporarily converge toward a common region in connectivity space. As training proceeds, the subtasks become more specialized again, and the context-dependent structure re-emerges.

In contrast, lazy networks show little reorganization of the relationships between subtasks. The similarity structure observed at initialization is largely preserved throughout training, with subtasks somewhat clustering by context. Thus, while rich networks transiently reorganize functional relationships among subtasks during learning, lazy networks largely preserve the initial subtask organization. This difference further highlights that learning in the rich regime involves substantial restructuring of the functionally relevant connectivity, whereas learning in the lazy regime primarily operates within the structure established at initialization.

Overall, these results extend previous findings from feedforward ANNs (*21*) to RNNs and network connectivity. While earlier work characterized qualitative differences between rich and lazy learning primarily in terms of function space, parameter magnitudes, and kernel analysis, our analyses complemented these insights with how these regimes diverge directly in the organization and evolution of the recurrent weight subspace in RNNs. Thus, the distinction between rich and lazy learning is reflected not only in performance and global parameter changes but also in how connectivity is structured and reshaped during training.

## Discussion

In this work, we aim to gain insights into how different functional modules are organized in a shared neural network connectivity and what factors are important to control this organization. Inspired by existing work, we systematically varied the variance of the recurrent weights at initialization and analyzed networks across multiple modeling choices (two activation functions, two training algorithms, and two neuroscience-inspired tasks). Consistent with findings in feedforward ANNs, we observed that lazy networks learn faster, as reflected in a more rapid decrease in training loss. However, when studying recurrent networks, other learning differences are not readily apparent in the full-rank recurrent weight matrices.

Instead, we were able to reveal crucial differences in network connectivity levels using the framework of operative dimensions. On the one hand, operative dimensions allow us to identify functionally relevant subspaces of the recurrent weight matrix for arbitrarily defined subtasks. Thereby, this framework enables going beyond global, functional descriptions of learning regimes on the level of network activities and instead analyzing how specific components of the tasks are implemented in the network’s recurrent connectivity. On the other hand, operative dimensions enabled us to isolate the functionally relevant components of connectivity while excluding weight subspaces that do not contribute to task execution.

Recent work similar to operative dimensions has examined these differences by analyzing the geometric properties of task-relevant manifolds extracted from network activity (*35*). Similar to the operative dimensions, this manifold-based approach focuses on network states actually visited during task execution, thereby isolating task-relevant components of the high-dimensional state space. This method is more tightly linked to the specific task definition, as distinct manifolds must be identified and separated for further analysis. In contrast, operative dimensions characterize the functionally relevant subspace of the recurrent connectivity and can be defined independently of an explicit task decomposition. Overall, operative dimensions proved essential for revealing important differences between the rich and lazy learning regimes.

### Differences across learning regimes

Most prominently, the subtask-specific operative dimensions become more shared with smaller variance in the initial weights (rich regime). In contrast, networks trained in the lazy regime implement different subtasks in more separated subspaces of the network connectivity.

Interestingly, we find a similar pattern in the subtask-specific subspaces of the network activity but less distinct across learning regimes. Analyses of low-rank RNNs have studied the relation between dominant activity dimensions and task-relevant directions in recurrent and input connectivity (*13, 23, 36*), and prior work has shown a correspondence between these two domains (*18*). In our setting, the observed alignment likely reflects that the network operates in a regime where variance generated by recurrent dynamics dominates input-driven or transient contributions. However, this similarity between activity- and connectivity-based subspaces cannot be expected in every network. The alignment between connectivity subspaces and network activity PCs depends on the relative contributions of recurrent, input-driven, and transient dynamics to the variance in the network activities.

Another qualitative difference between rich and lazy learning regimes shows that rich networks progressively adapt their operative subspace in the recurrent weight matrix **W** over training, as reflected by smaller subspace similarities (*S*) between training epochs and larger changes in operative dimensions as quantified by *AUE* and Δ*F*. Lazy networks, by contrast, learn primarily within the operative subspace defined at initialization. Interestingly, these results are also closely related to the distinction between *within-manifold* and *outside-manifold* learning described in neural population studies (*37*). In this framework, learning within an existing low-dimensional manifold enables rapid behavioral improvements without substantial reorganization of neural circuitry (lazy regime), whereas learning outside the manifold requires reshaping the underlying network dynamics and proceeds more slowly (rich regime).

### Mechanisms underlying distinct task organization

To understand the underlying mechanisms behind the qualitative differences in how rich and lazy networks organize subtasks in their network connectivity, we took a closer look at how the networks evolved during training. In particular, rich networks are characterized by more targeted weight updates (a larger fraction of the total recurrent weight change occurs within the operative subspace).

One plausible mechanism for these qualitative differences builds on our other observation showing that in rich networks, the input and output dimensions are more strongly aligned with the functionally relevant subspace of the recurrent connectivity. This becomes interesting in combination with previous analytical work. Specifically, Schuessler et al. (*30*) showed that gradient-based weight updates tend to be low-dimensional and aligned with input–output directions. Consistent with this, Hazelden et al. (*33, 34*) show that gradient-based weight updates are confined to the space spanned by the input and hidden activity at each update (self-referential bias). Note that Hazelden et al. operate on the space spanned by the network activities, whereas the operative dimensions operate directly on the recurrent weight matrix. The two levels of description are closely related but not identical, and we consider the insights gained to be complementary.

Taken together, these results suggest a mechanistic explanation: in rich networks, input and output dimensions are more closely aligned with the operative subspace of **W**, and since we are applying gradient-based update rules, the corresponding weight updates are more strongly aligned within the operative subspace (Fig. 5e). In contrast, in lazy networks, a much smaller proportion of weight changes occurs in the operative subspace of the recurrent connectivity. Hence, they have to achieve learning through changes in the input and output weights.

### Future work

An important question for future work is how these identified principles extend to more complex tasks requiring higher-dimensional computations. Previous theoretical work (*30, 33, 34*) has shown that gradient-based learning in neural networks is biased toward low-dimensional weight updates. While this property may be sufficient for the relatively structured tasks studied here, more complex tasks may require higher-dimensional solutions and richer computational representations. Implementing such solutions through inherently low-dimensional updates could become increasingly challenging. Thus, it remains unclear whether variations in the initial weight variance will nevertheless influence how different functional modules are organized within the same neural network when networks are required to support such more complex and high-dimensional computations.

Furthermore, many other commonly used hyperparameters are thought to influence how computations are organized within the network: networks using ReLU activations tend to form solutions in which functional submodules are more distinctly controlled by specific neuron groups, in contrast to activations that allow negative outputs (*2*). Similarly, the initialization of output weights has been shown to influence the nature of learned solutions. Schuessler et al. (*23*) demonstrated that initializing output weights with higher magnitudes leads to persistent task-irrelevant dynamics in the null space of the output vector, whereas smaller initial weights suppress such dynamics more effectively. How these factors interact with the rich/lazy axis identified here remains an open question.

### Conclusion

Our results show that the initial variance of the recurrent weight matrix determines how different tasks are organized in the recurrent connectivity. Conceptually, this is interesting because it demonstrates that distinct initial conditions can lead to qualitatively different learned task organizations, even when networks are trained with the exact same learning algorithm on the exact same task. Mechanistically, networks achieve this by aligning their input and output dimensions with the operative subspace of the recurrent weight matrix, thereby controlling the specificity of gradient-based weight updates.

For neuroscience, these insights are useful in two ways. First, they can guide modeling choices when using RNNs to model neural computation, since the choice of weight initialization is not a neutral implementation detail but a determinant of the resulting network’s functional organization.

Second, they offer a concrete, testable hypothesis for why some neural populations might predominantly exhibit feature-learning-like dynamics while others operate in a more kernel-like regime - for instance, if such populations differ systematically in the variability of their synaptic weights prior to or early in learning.

For machine learning, our results identify the initial variance of recurrent weights as a simple, practical hyperparameter for controlling the modularity of trained RNNs. This is relevant in any setting where the trade-off between shared and segregated representations matters, such as multitask learning, transfer learning, or continual learning, where overlapping subspaces can accelerate learning but may also increase interference between tasks.

## Acknowledgments

We would also like to thank Jean-Pascal Pfister for allowing us to use his computing power for part of the project.

## Author contributions

R.K. and V.M. designed the study and the methods. R.K. performed the analyses and wrote the manuscript with input from V.M.. R.K., V.M., reviewed the final manuscript.

## Data, Code, and Materials Availability

All analyses were run using MATLAB. We will provide all code and data required to reproduce the presented results here: https://github.com/ManteLab/operativeDimensions_taskOrganization For some colormaps, we used (*38*). No physical materials were generated in this work.

### Supplementary materials

Materials and Methods

Supplementary Text

Figs. S1 to S9

Tables S1

## References and Notes

1. S. Tafazoli, et al., Building compositional tasks with shared neural subspaces. Nature 650 (8100), 164–172 (2026), doi:10.1038/s41586-025-09805-2.

2. L. N. Driscoll, K. Shenoy, D. Sussillo, Flexible multitask computation in recurrent networks utilizes shared dynamical motifs. Nature Neuroscience 27 (7), 1349–1363 (2024).

3. V. Goudar, B. Peysakhovich, D. J. Freedman, E. A. Buffalo, X.-J. Wang, Schema formation in a neural population subspace underlies learning-to-learn in flexible sensorimotor problem-solving. Nature Neuroscience 26 (5), 879–890 (2023).

4. C. Tang, B. Lake, M. Jazayeri, Circuit explained: How does a transformer perform compositional generalization. PloS one 21 (2), e0340088 (2026).

5. R. Riveland, A. Pouget, Natural language instructions induce compositional generalization in networks of neurons. Nature Neuroscience 27 (5), 988–999 (2024).

6. J. J. Hopfield, Neural networks and physical systems with emergent collective computational abilities. Proceedings of the national academy of sciences 79 (8), 2554–2558 (1982).

7. R. Ben-Yishai, R. L. Bar-Or, H. Sompolinsky, Theory of orientation tuning in visual cortex. Proceedings of the National Academy of Sciences 92 (9), 3844–3848 (1995).

8. E. Gardner, The space of interactions in neural network models. Journal of physics A: Mathematical and general 21 (1), 257 (1988).

9. C. Van Vreeswijk, H. Sompolinsky, Chaos in neuronal networks with balanced excitatory and inhibitory activity. Science 274 (5293), 1724–1726 (1996).

10. H. Sompolinsky, A. Crisanti, H.-J. Sommers, Chaos in random neural networks. Physical review letters 61 (3), 259 (1988).

11. J. Aljadeff, M. Stern, T. Sharpee, Transition to chaos in random networks with cell-type-specific connectivity. Physical review letters 114 (8), 088101 (2015).

12. C. Curto, J. Geneson, K. Morrison, Fixed points of competitive threshold-linear networks. Neural computation 31 (1), 94–155 (2019).

13. F. Mastrogiuseppe, S. Ostojic, Linking connectivity, dynamics, and computations in low-rank recurrent neural networks. Neuron 99 (3), 609–623 (2018).

14. A. Dubreuil, A. Valente, M. Beiran, F. Mastrogiuseppe, S. Ostojic, Complementary roles of dimensionality and population structure in neural computations. biorxiv (2020).

15. A. Dubreuil, A. Valente, M. Beiran, F. Mastrogiuseppe, S. Ostojic, The role of population structure in computations through neural dynamics. Nature neuroscience 25 (6), 783–794 (2022).

16. A. Valente, J. W. Pillow, S. Ostojic, Extracting computational mechanisms from neural data using low-rank RNNs. Advances in Neural Information Processing Systems 35, 24072–24086 (2022).

17. F. Schuessler, A. Dubreuil, F. Mastrogiuseppe, S. Ostojic, O. Barak, Dynamics of random recurrent networks with correlated low-rank structure. Physical Review Research 2 (1), 013111 (2020).

18. R. Krause, M. Cook, S. Kollmorgen, V. Mante, G. Indiveri, Operative dimensions in unconstrained connectivity of recurrent neural networks. Advances in Neural Information Processing Systems 35, 17073–17085 (2022).

19. L. Chizat, E. Oyallon, F. Bach, On lazy training in differentiable programming. Advances in neural information processing systems 32 (2019).

20. T. Flesch, K. Juechems, T. Dumbalska, A. Saxe, C. Summerfield, Orthogonal representations for robust context-dependent task performance in brains and neural networks. Neuron 110 (7), 1258–1270 (2022).

21. M. Farrell, S. Recanatesi, E. Shea-Brown, From lazy to rich to exclusive task representations in neural networks and neural codes. Current Opinion in Neurobiology 83, 102780 (2023), 10.1016/j.conb.2023.102780, https://www.sciencedirect.com/science/article/pii/S0959438823001058.

22. A. Jacot, F. Gabriel, C. Hongler, Neural tangent kernel: Convergence and generalization in neural networks. Advances in neural information processing systems 31 (2018).

23. F. Schuessler, F. Mastrogiuseppe, S. Ostojic, O. Barak, Aligned and oblique dynamics in recurrent neural networks. Elife 13, RP93060 (2024).

24. P. J. Werbos, Generalization of backpropagation with application to a recurrent gas market model. Neural networks 1 (4), 339–356 (1988).

25. D. P. Kingma, J. Ba, Adam: A method for stochastic optimization. *arXiv preprint arXiv:1412.6980* (2014).

26. J. Martens, I. Sutskever, Learning recurrent neural networks with hessian-free optimization, in *ICML* (2011).

27. V. Mante, D. Sussillo, K. V. Shenoy, W. T. Newsome, Context-dependent computation by recurrent dynamics in prefrontal cortex. nature 503 (7474), 78–84 (2013).

28. D. Sussillo, O. Barak, Opening the black box: low-dimensional dynamics in high-dimensional recurrent neural networks. Neural computation 25 (3), 626–649 (2013).

29. N. Maheswaranathan, A. Williams, M. Golub, S. Ganguli, D. Sussillo, Universality and individuality in neural dynamics across large populations of recurrent networks. Advances in neural information processing systems 32 (2019).

30. F. Schuessler, F. Mastrogiuseppe, A. Dubreuil, S. Ostojic, O. Barak, The interplay between randomness and structure during learning in RNNs. Advances in neural information processing systems 33, 13352–13362 (2020).

31. G. R. Yang, M. R. Joglekar, H. F. Song, W. T. Newsome, X.-J. Wang, Task representations in neural networks trained to perform many cognitive tasks. Nature neuroscience 22 (2), 297–306 (2019).

32. Y. H. Liu, et al., How connectivity structure shapes rich and lazy learning in neural circuits. *arXiv preprint arXiv:2310.08513* (2023).

33. J. Hazelden, L. Driscoll, E. Shlizerman, E. Shea-Brown, KPFlow: An Operator Perspective on Dynamic Collapse Under Gradient Descent Training of Recurrent Networks. *arXiv preprint arXiv:2507.06381* (2025).

34. J. Hazelden, L. Driscoll, E. Shlizerman, E. Shea-Brown, The Global Empirical NTK: Self-Referential Bias and Dimensionality of Gradient Descent Learning. arXiv *preprint arXiv:2605.08746* (2026).

35. C.-N. Chou, H. Le, Y. Wang, S. Chung, Feature learning beyond the lazy-rich dichotomy: Insights from representational geometry. arXiv *preprint arXiv:2503.18114* (2025).

36. F. Mastrogiuseppe, J. Carmona, C. K. Machens, Stochastic activity in low-rank recurrent neural networks. PLOS Computational Biology 21 (8), e1013371 (2025).

37. J. A. Gallego, M. G. Perich, L. E. Miller, S. A. Solla, Neural manifolds for the control of movement. Neuron 94 (5), 978–984 (2017).

38. A. Biguri, Perceptually uniform colormaps, MATLAB Central File Exchange (2026), https://ch.mathworks.com/matlabcentral/fileexchange/51986-perceptually-uniform-colormaps.

