## Supplementary Materials for "Weight initialization shapes task organization in recurrent neural networks"

### Supplementary Materials for
### Weight initialization shapes task organization in recurrent neural networks

Renate Krauser*, Valerio Mante*

**This PDF file includes:**

Materials and Methods

Supplementary Text

Figures S1 to S9

Tables S1

#### Materials and Methods

##### Operative dimensions

The following section closely follows (*18*).

Consider an arbitrary dimension of the connectivity $\mathbf{a} \in \mathbb{R}^N$, we define $\hat{\mathbf{W}}$ as the matrix of rank $N-1$ obtained by removing the dimension $\mathbf{a}$ from the column space of $\mathbf{W}$ (for definition in row space see (*18*)):

$$\hat{\mathbf{W}} = \mathbf{W} - \mathbf{a}(\mathbf{a}^T\mathbf{W}) \tag{S1}$$

The dynamics of the resulting reduced-rank RNN are given by $\tau\dot{\hat{\mathbf{x}}}_t = -\mathbf{x}_t + \hat{\mathbf{W}}\mathbf{r}_t + \mathbf{B}\mathbf{u}_t + \boldsymbol{\sigma}_t$. At location $\mathbf{x}_t$ the activity evolves to $\hat{\mathbf{x}}_{t+1} = \mathbf{x}_t + \dot{\hat{\mathbf{x}}}_t dt$ over one time-step. The state of the full-rank network evolves to $\mathbf{x}_{t+1} = \mathbf{x}_t + \dot{\mathbf{x}}_t dt$ (for $\boldsymbol{\sigma}_t = 0$ in Eq. 1). We quantify the change in local dynamics due to removing dimension $\mathbf{a}$ in $\mathbf{W}$ as:

$$\Delta f = ||\mathbf{x}_{t+1} - \hat{\mathbf{x}}_{t+1}||_2 \tag{S2}$$

We then define the operative dimensions of $\mathbf{W}$ based on $\Delta f$ in two steps.

**Local operative dimensions.** In the first step, we infer a set of *local* operative dimensions at each of $P$ sampling locations $\mathbf{y}_j \in \mathbb{R}^N$ in state space. The sampling locations are chosen as an evenly distributed subset of the network states $\mathbf{x}_t$ that the network visits or passes nearby while performing the task. This choice of sampling locations ensures that the analysis focuses on local dynamics that are directly relevant to task performance (Fig. 2a; only a subset of $P > 100$ locations shown).

The *first* local operative dimension $\mathbf{d}_{1,j}$ at location $\mathbf{y}_j$ is defined as the dimension $\mathbf{a}$ that maximizes $\Delta f$ at location $\mathbf{y}_j$:

$$\mathbf{d}_{1,j} = argmax_{\mathbf{a}}(\Delta f)_{\{\mathbf{x}_t = \mathbf{y}_j\}} \tag{S3}$$

Up to $N-1$ further local operative dimensions $\mathbf{d}_{i,j}$ at the same location $\mathbf{y}_j$ are then defined in the same way, under the additional constraint to be orthogonal to all previously identified local operative dimensions at that location:

$$\mathbf{d}_{i,j} = argmax_{\mathbf{a}}(\Delta f)_{\{\mathbf{x}_t = \mathbf{y}_j\}}, \text{ constrained by: } \mathbf{d}_{i,j}^T\mathbf{d}_{i^*,j} = 0, \forall i^* < i \tag{S4}$$

Crucially, there exists an analytical solution to identify $\mathbf{d}_{i,j}$ ($\mathbf{d}_{1,j} = \mathbf{W}\mathbf{r}_j$ and $\Delta f = 0\ \forall i \geq 2$ (*18*)), which enables an efficient computation of the local operative dimensions even for a large number of sampling locations.

**Global operative dimensions.** In the second step, we define the *global* operative dimensions by combining the local operative dimensions $\mathbf{d}_{i,j}$ from all sampling locations $\mathbf{y}_j$ ($i = 1 : N$, $j = 1 : P$). Specifically, the local operative dimensions are scaled by their local $\Delta f$ and concatenated into one matrix $\mathbf{L}$ across all sampling locations $\mathbf{y}_j$.

$$\mathbf{L} = [\mathbf{d}_{1,1}\Delta f_{1,1}, \mathbf{d}_{1,2}\Delta f_{1,2}, \ldots, \mathbf{d}_{N,P-1}\Delta f_{N,P-1}, \mathbf{d}_{N,P}\Delta f_{N,P} \tag{S5}$$

where $\Delta f_{i,j} = \Delta f_{\{\mathbf{x}_t = \mathbf{y}_j, \mathbf{a} = \mathbf{d}_{i,j}\}}$. The i-th global operative dimensions $\mathbf{q}_i$ is then defined as the i-th left singular vector of $\mathbf{L}$:

$$\mathbf{L} = \Sigma_{i=1}^{N} \mathbf{q}_i g_i \mathbf{p}_i^T \tag{S6}$$

where $g_i$ are the singular values, and $\mathbf{p}_i$ the right singular vectors of $\mathbf{L}$. The global operative dimensions $\mathbf{q}_i$ consists of all the left singular vectors with $g_i$ being proportional to their functional importance. Note that defining global operative dimensions does not require the weight matrix to be constrained or parameterized in any specific way. This makes the framework applicable to networks trained or constructed using arbitrary methods. We refer to the global operative dimensions simply as the operative dimensions for brevity.

**Subtask-specific operative dimensions.** To compute these subtask-specific operative dimensions, it is sufficient to adjust the sampling locations $\mathbf{y}_j$. Instead of defining the sampling locations $\mathbf{y}_j$ as an evenly distributed subset of network states along trajectories across *all* input conditions, we restrict them to the states visited during a specific subtask. We then compute the local operative dimensions at these sampling locations and combine them to obtain the corresponding global operative dimensions as described in . More generally, operative dimensions can be defined for any arbitrary subset of sampling locations, enabling a flexible analysis of subtask-specific subspaces within the recurrent connectivity.

#### Subspace similarity

To quantify the similarity of the subspaces spanned by the subtask-specific operative dimensions, we use the concept of principal vectors: Let $\mathcal{U}, \mathcal{V} \subset \mathbb{R}^n$ be two linear subspaces of dimensions $d_u$ and $d_v$, respectively. The *principal vectors* $(u_i, v_i)$ are defined recursively as the pair of unit vectors that maximize the cosine similarity

$$\cos\theta_i = \max_{u\in\mathcal{U},\, v\in\mathcal{V}} \frac{u^\top v}{\|u\|_2\|v\|_2}, \tag{S7}$$

with the constraint that $(u_i, v_i)$ are orthogonal to all previously defined principal vectors. The corresponding angles $\theta_i \in [0, \pi/2]$ are the *principal angles* between the subspaces.

We define the similarity between two subspaces $\mathcal{U}$ and $\mathcal{V}$ as the average alignment of their first $k$ principal angles:

$$S(U,V) = \frac{1}{k}\sum_{i=1}^{k}(u_i^T v_i) \tag{S8}$$

where $k = \min(d_u, d_v)$ and $S_{U,V} \in [0,1]$. This measure is bounded between 0 and 1, with $S(\mathcal{U},\mathcal{V}) = 1$ if and only if the subspaces are identical, and $S(\mathcal{U},\mathcal{V}) = 0$ if they are orthogonal. Notably, it is not straightforward to define a principled threshold for the number of operative dimensions that constitute the functionally relevant weight subspace or to assess the similarity of subspaces with different dimensionalities. To avoid such ambiguities, we restrict our comparison to the subspace spanned by the first $k = 5$ operative dimensions between network types.

#### Relation between targeted weight updates and operative subspace

It is not entirely trivial how changes to the weights within the operative subspace relate to changes in the operative subspace itself. To build an intuition, consider a single location in the state space. At such a point, analytical results show that the operative subspace is one-dimensional (*18*). If a weight update is performed exactly along this dimension (i.e., a targeted update fully within the local operative subspace), it will modify the weights within the subspace without changing the subspace itself. In contrast, a weight update along a direction outside the operative subspace will shift the operative subspace.

However, this intuition does not directly extend to the global operative subspace defined across multiple sampling locations. The global operative dimensions are constructed as an *average* over local operative dimensions (Eq. S6). As a result, even weight updates that lie within the global operative subspace can alter it. A direction that belongs to the global operative subspace may not align perfectly with the local operative subspace at a given state, and thus corresponds locally to a component outside the operative subspace. Consequently, updates within the global operative subspace can affect the overall operative subspace alignment. Nevertheless, more relevant global operative dimensions tend to be better aligned with a larger fraction of local operative dimensions, thereby inducing smaller shifts in the global subspace than updates along less relevant dimensions.

Hence, the observed pattern in the rich learning regime with more weight updates within the operative subspace while simultaneously also changing the operative subspace is counterintuitive at first. More targeted weight updates should lead to smaller changes to the operative subspace overall. However, one has to keep in mind that rich networks generally yield lower-dimensional operative subspaces. Hence, global operative dimensions of the same rank are less functionally relevant in rich networks than in lazy networks and consequently lead to larger changes in the subspace alignment in rich than in lazy networks.

#### Supplementary Text

##### Network training

The training was not equally stable across different network settings, tasks, activation functions, optimization methods, and weight initializations, but all used networks show a performance clearly above chance level. To ensure a fair comparison across these network families, we trained between 10 and 30 networks per setting (Tab. S1). For the subsequent analyses, we restricted our dataset to the nine best-performing networks within each family, based on the minimal loss (Eq. 3). An exception occurred for the pSWG, tanh, HF configuration. In this case, we trained 50 networks, but only four converged to valid solutions with a low-variance weight initialization (0.0001). Consequently, only four networks were included in the analysis for this family, whereas nine networks were used for all other families.

##### Dimensionality of recurrent weight matrix

To confirm that the recurrent weight matrices remain high-dimensional despite the existence of low-dimensional functionally relevant subspaces, we analyzed the dimensionality of the trained recurrent connectivity matrices $\mathbf{W}$. As shown in Fig. AS1, the full recurrent weight matrices retain a high dimensionality across all network families and learning regimes. Thus, although operative dimensions reveal that only a small subset of the connectivity is required to reproduce the learned network function, the underlying recurrent weight matrices are not themselves low-dimensional. This highlights the distinction between the dimensionality of the full connectivity and the dimensionality of the functionally relevant subspace used by the network.

##### Dimensionality of operative dimensions and recurrent weights over training

Here, we analyze how the dimensionality of the operative dimensions evolves over the course of training (Fig. AS2). As a result, we find distinct patterns across learning regimes. In rich networks, the dimensionality typically starts relatively high, rapidly decreases during early training, and then stabilizes at an intermediate level as learning converges. In contrast, lazy networks show little change in dimensionality over training, with the functional subspace remaining comparatively stable throughout. These results further highlight that rich learning involves a substantial reorganization of the recurrent connectivity, whereas lazy learning relies on a more structurally stable functional subspace.

##### Similarity of subtask-specific operative dimensions

To test whether the organization of subtask-specific operative dimensions generalizes across tasks, we performed the same analysis on the phase-dependent sine-wave generator (pSWG) task, as shown in the main text for the context-dependent integration task (Fig. 3e-h). Similar to the CTXT task, we compared how well operative dimensions derived from one subtask reproduced the dynamics of another by measuring the resulting $S$ (Fig. AS3a,b) and $AUE$ (Fig. AS3c,d) across subtasks. We again find that each subtask is implemented in more shared subspaces for rich networks than for lazy networks. Thus, the structured organization of subtask-specific operative dimensions is not unique to the CTXT task but generalizes across different task structures and network dynamics.

##### Similarity of subtask-specific operative dimensions across training

To analyze how the relationship between subtask-specific operative dimensions develops during training, we evaluated the $AUE$ across subtasks over the course of training. For each network and each training iteration, we computed the $AUE$ for every pair of subtasks by applying the operative dimensions obtained from one subtask to the weight matrix of all other subtasks (see also Fig. 2d). We then averaged these values across all subtask pairs at each training iteration.

At initialization (training iteration 0), we found that rich networks exhibit a substantially higher average $AUE$ across subtasks compared to lazy networks (Fig. AS4a). This indicates that prior to training, the subtask-specific operative dimensions in rich networks are more distinct from one another, whereas lazy networks start with more similar subtask subspaces.

Next, we examined how this average $AUE$ changes over training. We computed the average change in $AUE$ ($\Delta AUE$) separately for the early training phase (training iteration 1–20, Fig. AS4b) and the later phase (training iteration 21–120, Fig. AS4b). During the initial phase, rich networks show a clear decrease in average $AUE$, indicating that their subtask-specific subspaces become more similar as training progresses. In contrast, lazy networks show very little change in this period. During the later training phase, the average $AUE$ remains relatively stable in both regimes, suggesting that the similarity structure between subtask subspaces is largely fixed after the initial learning stage.

Together, these results indicate that training actively pushes subtask subspaces toward one another in rich networks, primarily during the early phase of learning, whereas lazy networks preserve the similarity structure present at initialization. This confirms our results presented in Fig. 6.

##### Similarity of subtask-specific activity subspaces

We also asked whether the structured organization observed in the operative dimensions is also reflected in the network activity of the pSWG task. As in the main text for the CTXT task (Fig. 4d-e), we compared the similarity of subtask-specific activity subspaces by computing the overlap between the first principal components of network activity across subtasks.

Consistent with the results for the CTXT task, we find that activity subspaces in the pSWG task are also only partially shared across subtasks, while still exhibiting a structured organization related to the underlying task conditions (Fig. AS5). Furthermore, the same trend is observed across learning regimes: rich networks show more overlapping activity subspaces, whereas lazy networks exhibit increasingly separated subtask-specific activity representations. These findings demonstrate that the relationship between learning regime and subtask organization generalizes across connectivity and activity space, as well as across different task settings.

##### Correlation between similarity measures

The choice of the number of dimensions used in the similarity analyses is, in principle, an arbitrary parameter; in the main text, we fixed this value to $n = 5$. Here we show that this choice is not critical: for $n \geq 4$, the qualitative relationship between $AUE$ and subspace similarity $S$ remains stable, and the correlation between these two measures is preserved (Fig. AS6).

Specifically, to assess whether our conclusions depend on the number of operative dimensions used to define functionally relevant subspaces, we conducted additional control analyses by varying this parameter. Specifically, we computed the area under the error curve ($AUE$) obtained by applying operative dimensions extracted from the untrained network to the fully trained network and vice versa. In parallel, we quantified the subspace similarity $S$ between operative dimensions of the untrained and fully trained networks while systematically varying the number of dimensions used to construct the subspaces.

Note that directly comparing the similarity between subtask-specific operative subspaces should be interpreted with caution for two reasons. First, the subspaces may differ in dimensionality, introducing bias because higher-dimensional subspaces are more likely to overlap. Second, the operative dimensions are ordered by functional relevance; comparing plain subspaces ignores this ordering and therefore discards information about the relative importance of individual dimensions. Nevertheless, our analyses show that the computed subspace similarities are largely consistent with the $AUE$ measurements (Fig. AS6).

##### Changes in network weights and functional subspaces over full training

For completeness to characterize how learning differs between the rich and lazy regimes, we quantified the total changes in recurrent, input, and output weights across the full training procedure

(Fig. AS7a–c). While lazy networks tend to exhibit somewhat larger changes in input and output weights, the total changes in the recurrent connectivity $\mathbf{W}$ are overall comparable across learning regimes. Thus, differences between rich and lazy learning are not fully captured by the magnitude of recurrent weight changes alone. We therefore additionally examined how strongly the functionally relevant recurrent subspaces change over training. To this end, we computed the $AUE$ between operative dimensions extracted from untrained and fully trained networks (Fig. AS7d–e). Consistent with the analyses in the main text, rich networks show larger $AUE$ values, indicating that the operative dimensions before and after training are less interchangeable than in lazy networks. Hence, although overall recurrent weight changes are similar across regimes, the functionally relevant recurrent subspace is reorganized more strongly in rich networks during learning.

Additionally, we examined how the standard deviation of the recurrent weight matrix $\mathbf{W}$ evolves throughout training (Fig. AS8d–e). In lazy networks, the variance increases only slightly over the course of learning, whereas rich networks exhibit a substantially larger increase in weight variance. This difference further explains why the distinction between the rich and lazy learning regimes is most pronounced during the early stages of training, when the initial difference in weight variance is largest. As training progresses, the recurrent weight distributions of the two regimes become more similar, reducing this distinction.

**Robustness of qualitative differences across learning regimes**

To assess the robustness of our results (Fig. 5a and f) with respect to the number of operative dimensions used to define the functionally relevant subspace, we repeated the analysis of targeted weight updates and similarity of consecutive operative subspaces using different numbers of operative dimensions (Fig. 5). Across all network types, we find that the results are qualitatively consistent for different numbers of operative dimensions ($n \geq 3$). While the absolute values vary slightly with the dimensionality choice, the relative differences between rich and lazy learning regimes remain qualitatively unchanged in that the rich regime performs more targeted weight updates (Fig. AS9a) and larger changes in the used operative subspaces (Fig. AS9b). In the main text, we therefore report results using five operative dimensions as a representative and stable choice.

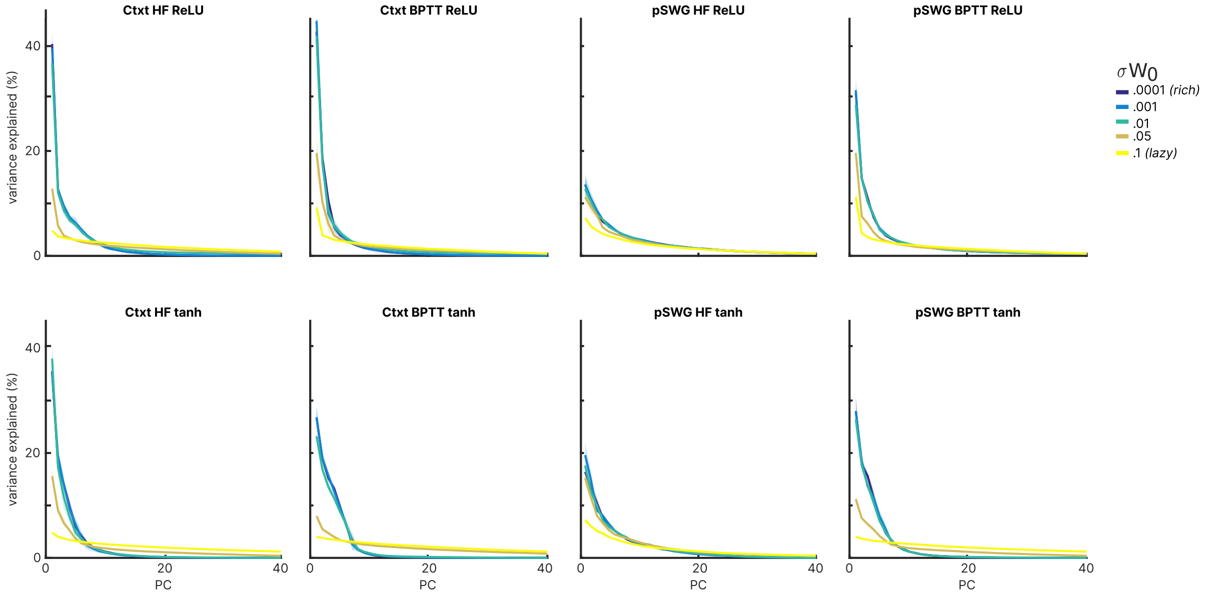


**Figure S1**: **Dimensionality of recurrent weight matrix W after training.** Shaded area: $mad$ over all networks per $\sigma W_0$.

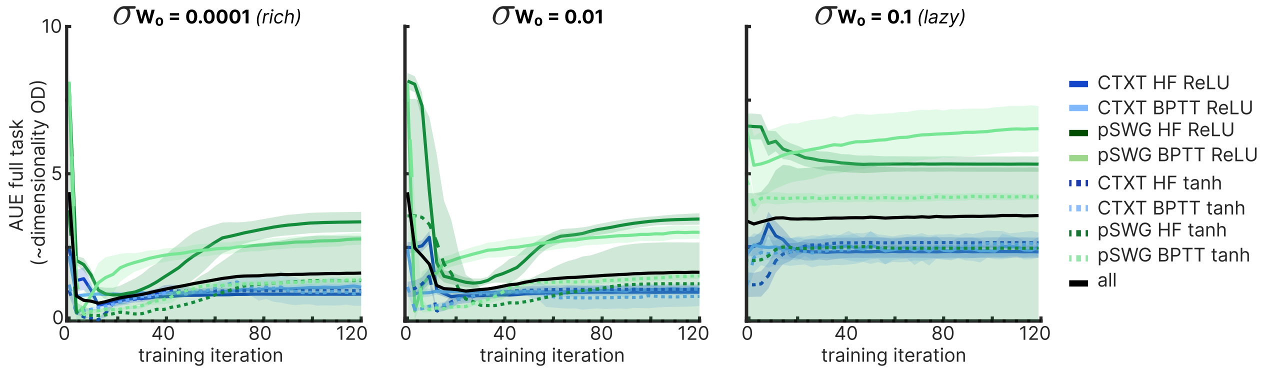


**Figure S2**: **Dimensionality of operative dimensions over training** $AUE$ of full task over training iterations for all network families and three different weight initialization values. Shaded area: $mad$ over all networks per $\sigma W_0$.

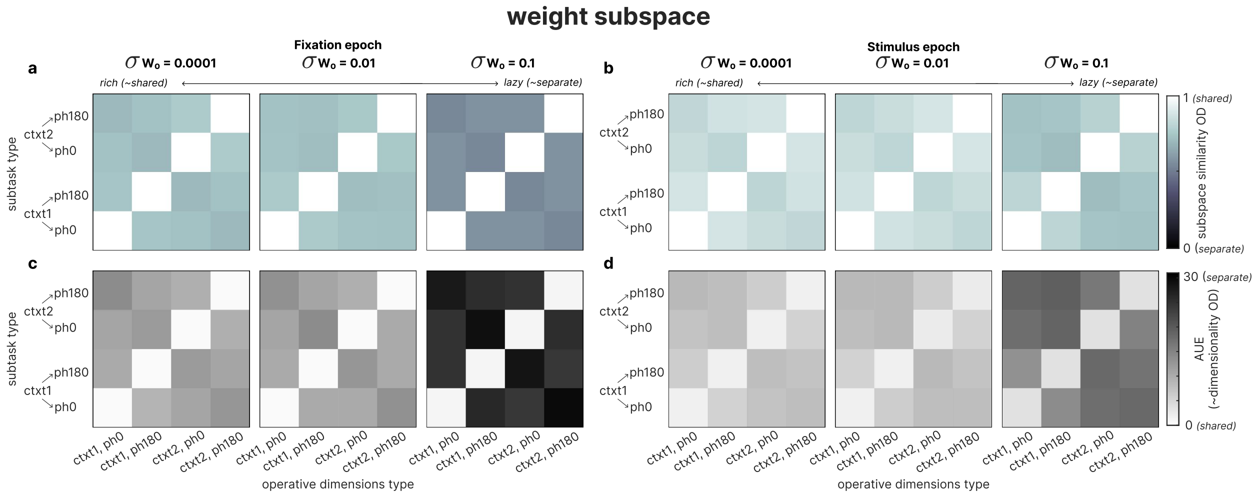


**Figure S3**: **Task representations in network connectivities (pSWG).** (**a**) Pairwise similarity of subtask-specific operative dimensions of fixation period in pSWG (averaged over networks) over varying weight initializations $\sigma W_0$. (**b**) same as (**a**) for stimulus period. (**c**) $AUE$ of subtask-specific operative dimensions of fixation period in pSWG (averaged over networks) over varying weight initializations $\sigma W_0$. $AUE$ is calculated for each subtask type (y-axis) using all subtask-specific operative dimension types (x-axis). (**d**) same as (**c**) for stimulus period.

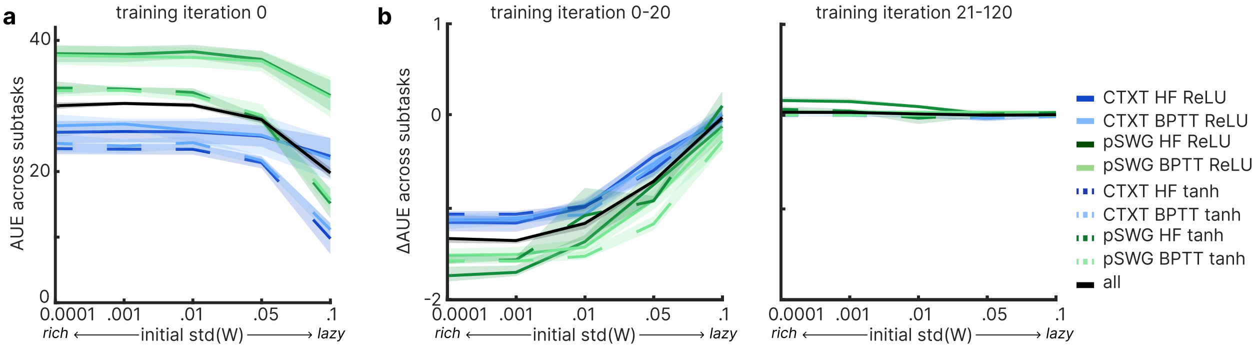


**Figure S4**: **Similarity of subtask-specific operative dimensions across training.** (**a**) Average $AUE$ across all subtasks tested with (unmatching) operative dimensions of all other subtasks at weight initialization (averaged over networks) over varying weight initializations $\sigma W_0$. (**b**) Average change in $AUE$ across all subtasks tested with (unmatching) operative dimensions of all other subtasks during iteration 1-20 and 21-120 (averaged over networks) over varying weight initializations $\sigma W_0$. (**a**)-(**b**) shaded area: median absolute deviation ($mad$) over all networks per $\sigma W_0$.

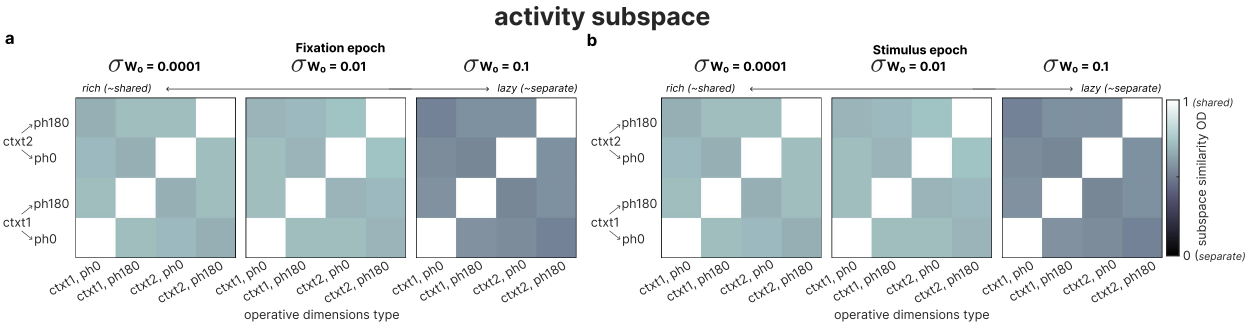


**Figure S5**: **Task representations in network activities (pSWG).** (**a**) Pairwise similarity of subtask-specific network activity subspaces of fixation period in pSWG (averaged over networks) over varying weight initializations $\sigma W_0$. (**b**) same as (**a**) for stimulus period.

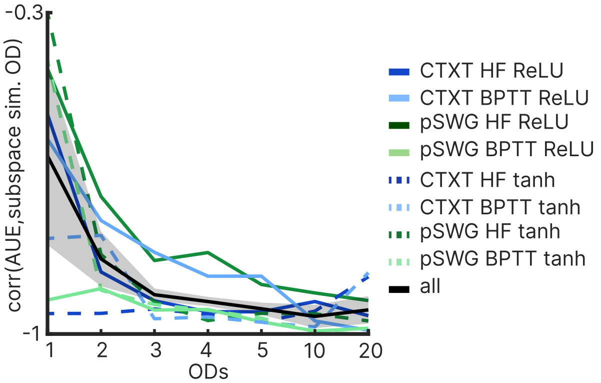


**Figure S6**: **Correlation between similarity measures over subspace dimensionality.** Correlation between $AUE$ and $S$ between operative dimensions of untrained and trained networks, shown for different numbers of operative dimensions. The correlation is stable for $n \geq 3$, indicating robustness to the choice of dimensionality. Shaded area: median absolute deviation ($mad$) over all networks per $\sigma W_0$.

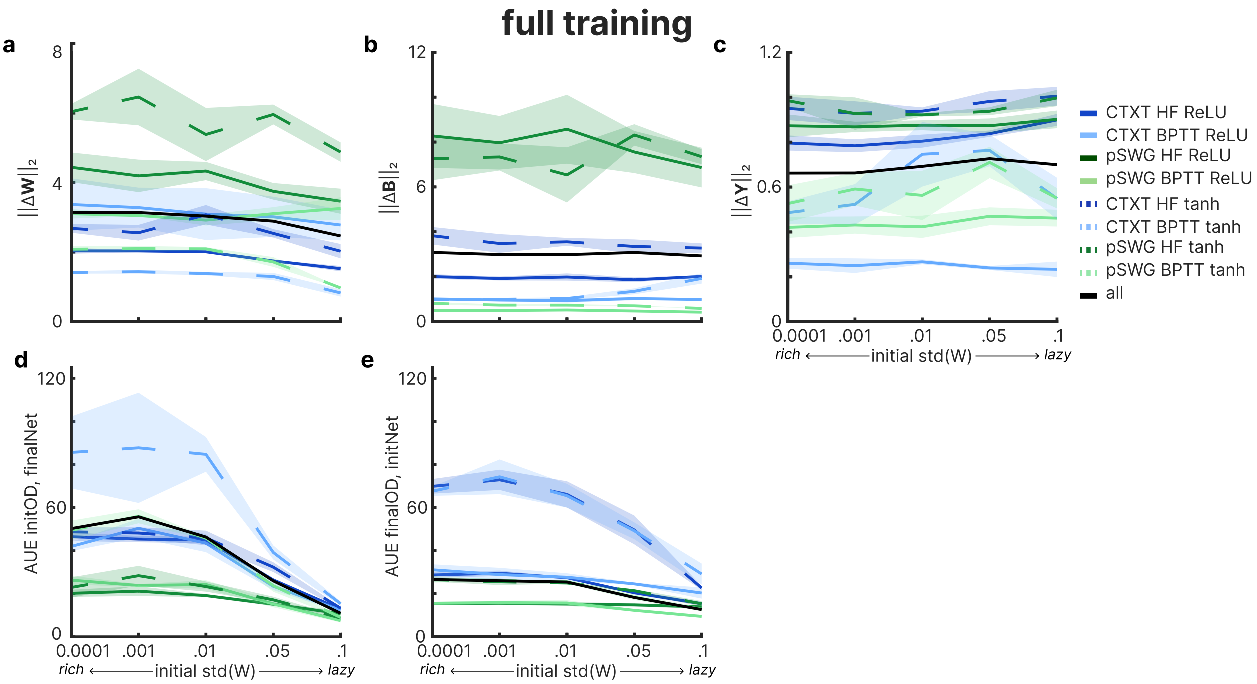


**Figure S7**: **Network change over training.** ( (**a**) Total difference (L2-norm) in recurrent weights **W** over training. (**b**) Total difference (L2-norm) in input weights **B** over training. (**c**) Total difference (L2-norm) in output weights **Y** over training. (**d**/) $AUE$ for operative dimensions of the untrained network applied to the network after training. (**e**/) $AUE$ for operative dimensions of the fully trained network applied to the untrained network. (**a**)-(**e**) shaded area: median absolute deviation ($mad$) over all networks per $\sigma W_0$.

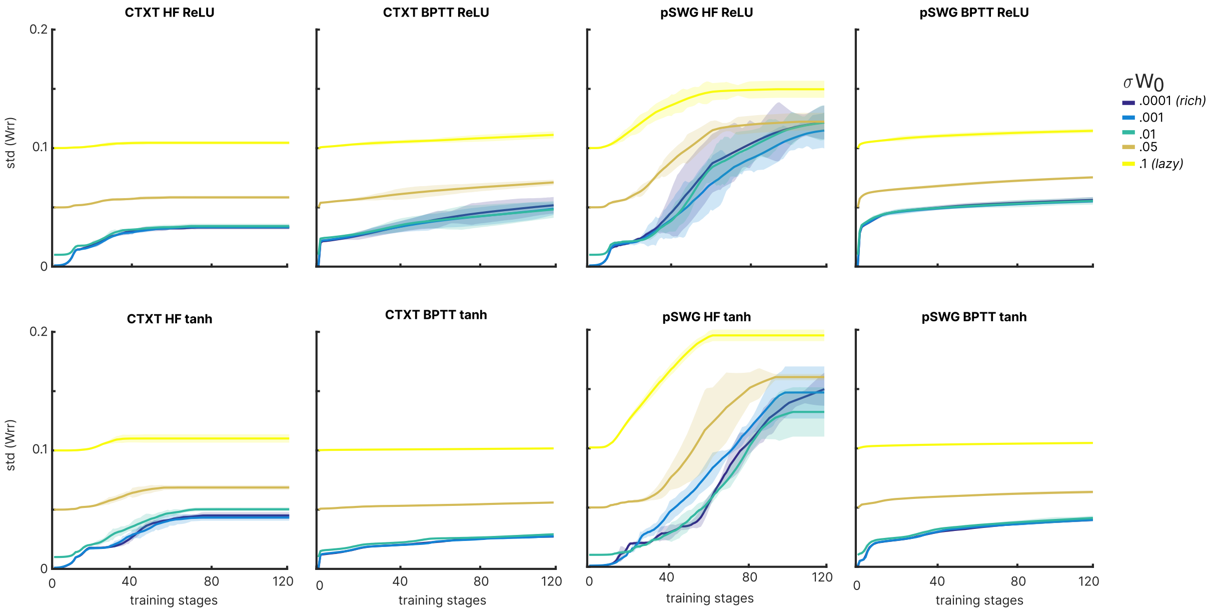


**Figure S8**: **Standard deviation of recurrent weight matrix W over training.** Shaded area: median absolute deviation ($mad$) over all networks per $\sigma W_0$.

| $\sigma W_0$ | **0** | **0.001** | **0.01** | **0.05** | **0.1** |
|---|---|---|---|---|---|
| **CTXT HF relu** | 10 | 10 | 10 | 10 | 10 |
| **CTXT BPTT relu** | 30 | 30 | 30 | 10 | 30 |
| **pSWG HF relu** | 10 | 10 | 10 | 10 | 10 |
| **pSWG BPTT relu** | 10 | 10 | 10 | 10 | 10 |
| **CTXT HF tanh** | 10 | 10 | 10 | 10 | 10 |
| **CTXT BPTT tanh** | 30 | 30 | 30 | 10 | 10 |
| **pSWG HF tanh** | 50 | 50 | 50 | 50 | 50 |
| **pSWG BPTT tanh** | 30 | 30 | 30 | 30 | 30 |

**Table S1**: **Number of trained networks per network family.** CTXT: context-dependent integration; pSWG: phase-dependent sine wave generation; HF: Hessian-free optimization; BPTT: Backpropagation through time; relu: Rectified Linear Unit activation function; tanh: tanh-activation function

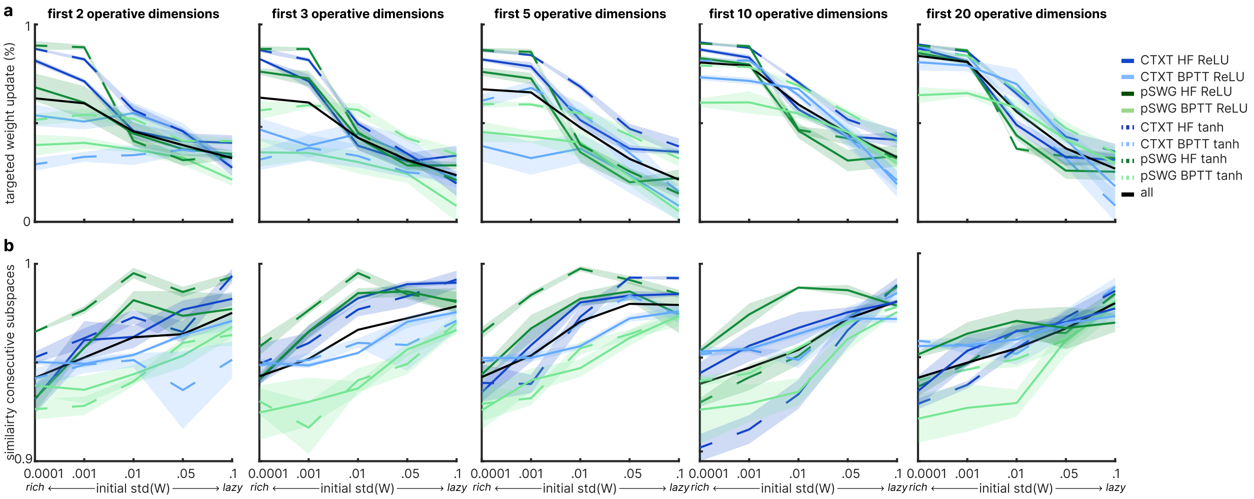


**Figure S9**: **Qualitative learning differences across subspace dimensionality** (**a**) Proportion of targeted recurrent weight updates within the functionally relevant subspace, computed using different numbers of operative dimensions (averaged over training iterations 1-20). (**b**) Average similarity of consecutive operative subspaces, computed using different numbers of operative dimensions (averaged over training iterations 1-20). (**a-b**) Results are shown for subspaces spanned by the first n operative dimensions ($n = 2, 3, 5, 10, 20$). Shaded area: median absolute deviation ($mad$) over all networks per $\sigma W_0$

.
